# TedSim: temporal dynamics simulation of single cell RNA-sequencing data and cell division history

**DOI:** 10.1101/2021.06.21.449283

**Authors:** Xinhai Pan, Hechen Li, Xiuwei Zhang

## Abstract

Recently, the combined scRNA-seq and CRISPR/Cas9 genome editing technologies have enabled simultaneous readouts of gene expressions and lineage barcodes, which allows for the reconstruction of the cell division tree, and makes it possible to trace the origin of each cell type. Computational methods are emerging to take advantage of the jointly profiled scRNA-seq and lineage barcode data to better reconstruct the cell division history or to infer the cell state trajectories.

Here, we present TedSim (single cell **Te**mporal **d**ynamics **Sim**ulator), a simulator that simulates the cell division events from the root cell to present-day cells, simultaneously generating the lineage barcodes and scRNA-seq data. In particular, TedSim generates cells from multiple cell types through cell division events. TedSim can be used to benchmark and investigate computational methods which use either or both of the two types of data, scRNA-seq and lineage barcodes, to study cell lineages, ancestral cells or cell trajectories. TedSim is available at: https://github.com/Galaxeee/TedSim.

## Introduction

Understanding how single cells divide and differentiate into different cell types in developed organs is one of the major tasks of developmental biology. The temporal dynamics of cells are being studied by various approaches from different aspects. First, *trajectory inference* (TI) methods [1–5] are developed to infer the trajectories of cells from single cell gene expression data, typically obtained from single cell RNA sequencing (scRNA-seq) experiments. Trajectory inference methods make the assumption that although the cells sequenced together are collected at the same time, they represent different stages of the temporal dynamic process in the cell populations. Prevalent trajectory inference methods aim to find the trajectory backbone which represents the major cell states and the dynamic paths between the states, and then “sort” the cells onto the backbone structure. These methods have assisted biological discoveries in various biological systems [6–9], however, it is not clear whether using only the scRNA-seq data of present-day cells can always reconstruct the cell trajectories, as some information may have been lost during the developmental processes and not captured in the scRNA-seq data [10–12].

Second, *lineage tracing* technologies using CRISPR/Cas9 genome editing technology blaze a new trail to study the cell developmental mechanisms. A barcode is inserted into the genome which accumulates CRISPR-induced mutations during the process of the cell divisions, and the mutated barcode can be read out together with the gene expression profile in a cell through scRNA-seq [10,13–15]. This barcode will be referred to as *lineage barcode* in this manuscript. Thus, the output of these lineage tracing experiments is composed of two types of data: the single cell gene expression and the lineage barcode of each cell. Computational tools to reconstruct the cell division history of single cells from the lineage barcodes are widely explored [16, 17]. In an ideal scenario, the reconstructed cell division tree along with the gene expression profiles of present-day cells can provide insights into how different cell types originate from progenitor cells, and allow us to draw the cell fate map at an unprecedented high resolution. However, the missing data in the lineage barcodes and the large cell amount pose challenges for cell lineage tree reconstruction methods, resulting in low-resolution, inaccurate reconstructed trees.

With simultaneously profiled single cell gene expression data and lineage barcodes, methods that integrate the two modalities start to emerge, aiming to achieve better results than the unimodal methods in terms of inferring cell lineage and developmental trajectories. However, few methods have been developed and their performances are unclear [17, 18].

Previously, computational tools to simulate either single cell gene expression data [19–21] or lineage barcode data [22] were proposed. These simulation tools can provide quantitative evaluations of existing computational methods and set baselines for the development of new methods. The single-cell gene expression simulators mostly sample gene expression levels of single cells from probability distributions of the data under certain statistical assumptions [19–21], i.e. the cells of the same cell type are i.i.d. samples centered at pre-calculated mean expressions of the cell type. The synthetic expressions do not maintain the actual lineage relationships between single cells. On the other hand, Salvador-Martínez *et al* developed a simulator to generate the lineage barcode information by simulating the cell division processes but not the gene expression data [22].

The joint profiling of single cell gene expression and the lineage tracing barcode data allow for the analysis of the origin of various cell types. Previously, the lineage barcodes and the gene expression data were utilized to reconstruct the lineage tree and identify the cell types separately [10,13–15,23]. Recently, hybrid methods that attempt to use both types of information are being designed to either use the gene expression information to obtain better lineage trees [17] or to use the lineage information to obtain better inferred trajectories from gene expression data [18]. To study the relationship between gene expression and lineage origins and to benchmark the hybrid methods, we need to generate both single cell gene expression and lineage barcode information simultaneously along the cell division processes with ground truth information on the cell division history and cell state transition.

In this paper, we present TedSim (single cell **Te**mporal **d**ynamics **Sim**ulator), the first simulator that outputs combined readouts of single cell gene expression and lineage barcode data. Unlike existing simulators for scRNA-seq data [19,21,24], TedSim simulates the actual cell division events, and obtains cells of different cell types at different stages of the developmental trajectories, alongside with lineage barcodes through genetic scarring during cell divisions.

In order to simulate the accumulation of CRISPR/Cas9 genetic scarring, it is necessary to iterate along generations of cell divisions to model the stochastic occurrence of mutation events. One challenge of designing TedSim is how to obtain various cell types through cell divisions. We present a simple model that utilizes *asymmetric cell divisions* [25–27] to assign cell types on the cell division tree. We consider an underlying *cell state tree*, which represents the developmental relationships of cell states, as the input to TedSim. The cell state tree will help to determine the states of the cells on the cell division tree (Fig. 1). The state shifts of cell identities during cell divisions are achieved by asymmetric cell divisions, and at the same time genetic scarring events can happen and be inherited by the daughter cells. Identities of cells are represented using latent space vectors which are determined by both their specific cell type (determined by the state tree) and the identities of the ancestors on the lineage tree. The gene expression profile of each cell is then generated from the cell identity of the cell.

**Figure 1.**
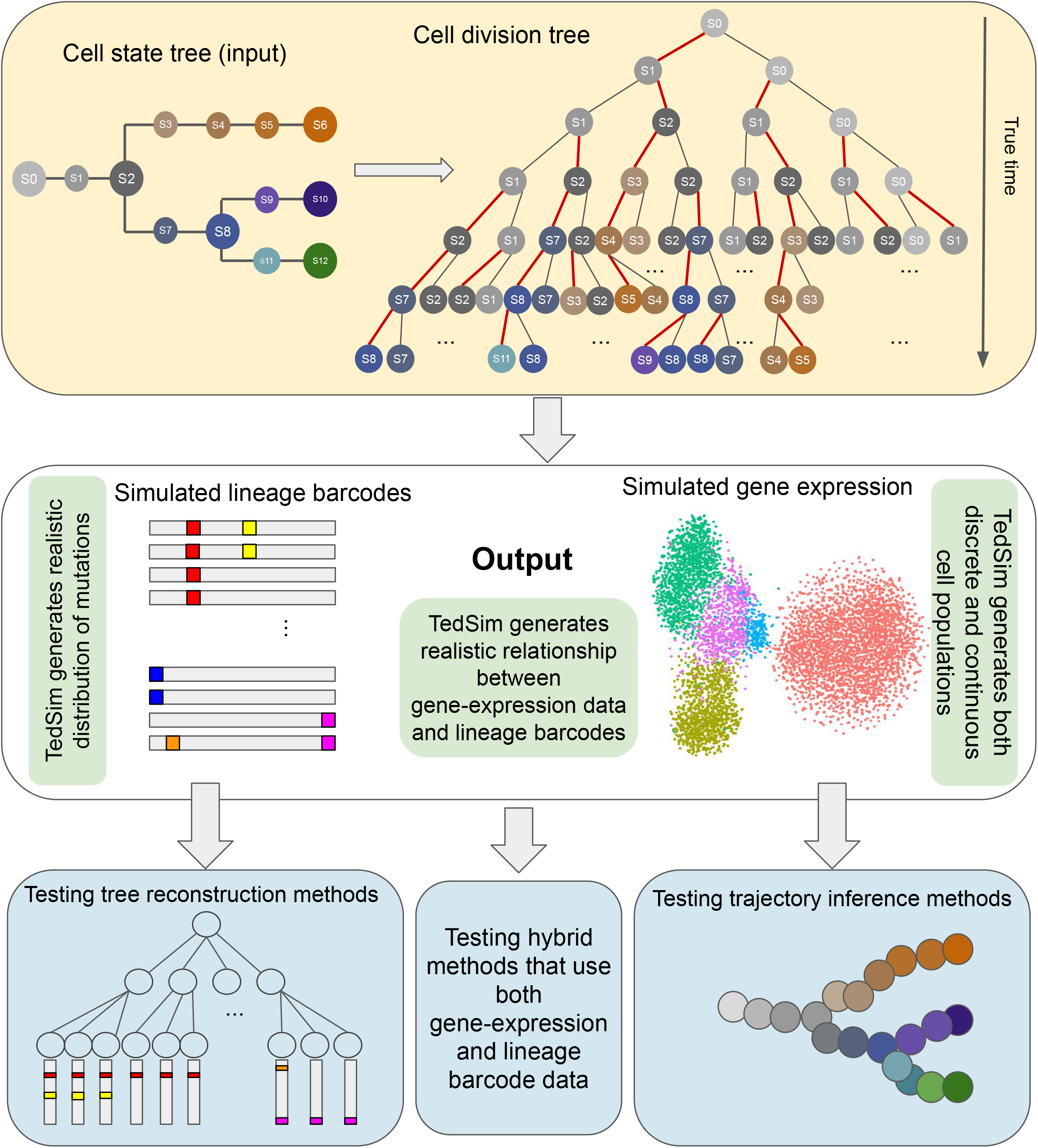
TedSim workflow. The cell state tree models the true trajectories of cell differentiation and the cell division tree represents the true lineage. By traversing the cell division tree from root to leaves, the cell states are determined based on the state tree and the asymmetric division events (denoted by red edges in the cell division tree). TedSim simultaneously simulates gene expressions and lineage barcodes of all the cells, while the state and lineage relationships of the cells are maintained. The simulated data can be used to test not only trajectory inference or tree reconstruction methods, but also hybrid methods which use both gene expression and lineage barcode data.

We show that TedSim can generate both discrete and continuous cell populations, lineage barcode data with similar mutation distribution as observed in real data, and in particular, realistic “relationship” between the lineage barcode and gene expression data (Fig. 1). By “relationship” we particularly look into the following question: do cells from the same cell types (defined by their gene expression profiles) necessarily originate from the same clone? We show that similar to that in real data, in our simulated data, each clone can have multiple cell types and one cell type can spread over multiple clones.

We demonstrate the application of TedSim in benchmarking the following computational methods: 1) With synthetic lineage barcodes of single cells, we can benchmark lineage tree reconstruction methods; 2) With simulated scRNA-seq data, we can apply the trajectory inference methods and test whether using only the present-day cells can recover the underlying cell state tree; 3) With both lineage barcode and scRNA-seq data, we can benchmark methods which combine gene expression and lineage barcodes. In particular, we discuss the limitations of current methods which integrate the two types of data.

## Results

### TedSim generates both continuous and discrete populations through cell divisions

That TedSim simulates scRNA-seq data through actual cell divisions is a key feature which distinguishes TedSim from the state-of-the-art simulators for scRNA-seq data. In this way, TedSim directly models the temporal dynamics of single cells and thus can be used to analyze the underlying relationships of lineage and gene expressions. By modeling cell divisions which are basic events that lead to various cell types, TedSim can naturally generate discrete or continuous populations in the present-day cells, with the same underlying cell state transition mechanisms.

In order to obtain cells with diverse cell types through cell divisions, we adopt the fundamental concept of *asymmetric divisions*. In biology, stem cells are characterized by their ability to self-renew and produce differentiated phylogeny which is achieved through controlled *asymmetric divisions* [25–27]. Asymmetric division is a key process for a cell to divide into two cells with different cellular fates, where one cell remains at the same state as the parent, and the other shifts to the future state that will eventually develop into fully-differentiated cells. TedSim utilizes a *cell state tree* which models the cell differentiation process to determine the future state when a cell divides asymmetrically. Starting from the root of the cell lineage tree, a cell can divide either symmetrically into two cells of the same state as their parent, or asymmetrically as described above. By incorporating the parent cells’ identity when generating the identity of the daughter cells, we are able to combine heterogeneities from both its cell state and lineage path. Transcriptomic data simulated by TedSim can reflect the true trajectory as demonstrated in the cell state tree while being compounded by lineage relationships between cells, which reflects the scenario in reality [28].

The process of how TedSim simulates scRNA-seq data is shown in Fig. 2a. The cell state tree and cell lineage tree (Fig. 2b) provide two different sources of heterogeneities in the gene expression profiles of cells, where the former represents the underlying programmed cell fate decision mechanism, and the latter represents effects from clonal origins of cells.

**Figure 2.**
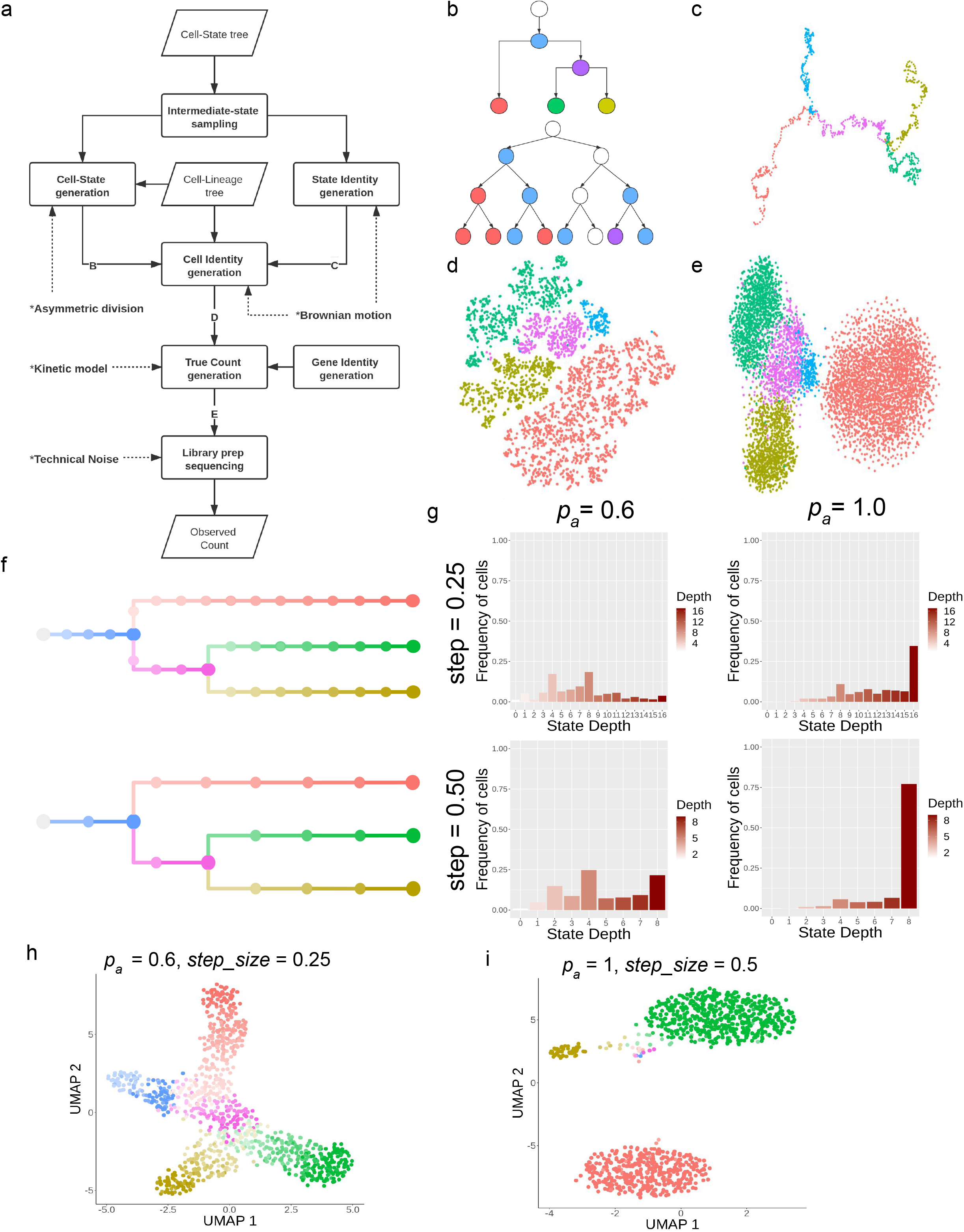
Simulation of scRNA-seq data in TedSim. (a) Flowchart of generating observed gene expression counts. Parallelogram represents input/output and rectangle represents process. (b-e) Key intermediate data corresponding to (b-e) in Figure 2a. (b) Cell state tree and cell lineage tree where each cell is colored with its state. Asymmetric division is applied so that the state shift happens to only one of the daughter cells at each division. (c) 2-d PCA visualization of simulated State Identity Vectors (SIVs) along the cell state tree. The length of the SIVs is 30 (20 diff-SIVs and 10 non diff-SIVs). (d) 2-d tSNE visualization of Cell Identity Vectors (CIVs) of 2048 cells. A CIV has the same length as a SIV. (e) 2-d tSNE visualization of true counts of 2048 cells and 500 genes. (f) Cell state trees sampled with two different *step_size =* 0.25, 0.5, colors represent cell states in (h) and (i). (g) The frequency of cells belonging to each cell state in the simulated dataset, stratified by the depth of the corresponding state in the cell state tree. (h) 2-d UMAP visualization of a continuous population of cells where *p_a_* and *step_size* are small, and the trajectory of the states can be observed. (i) 2-d UMAP visualization of a discrete population of cells where *p_a_* and *step_size* are large, and the cells are separated into discrete clusters.

We assume that cell states change gradually along the given cell state tree, and each state is represented by a state identity vector (SIV) which can be considered as the latent representation of the state. We generate the SIVs along the cell state tree from the root using the Brownian motion model which was used to model the evolution of genetic traits [29]. The changes of trait values of the Brownian motion are drawn from normal distributions, and the covariance matrix of the trait values corresponds to the given tree structure (Methods, Supplementary Note Sec. 3). Fig. 2c shows the 2-d PCA visualization of the SIVs generated along the cell state tree in Fig. 2b. To obtain a desired number of cell types in simulated data, we sample states from the cell state tree by taking the SIVs at both ends of each edge and some intermediate positions of each edge where each sampled state corresponds to a cell type. For example, in the cell state tree shown in Fig. 1, we take states S_0_, S_1_, …, S_12_ as the discretized states to guide the state transitions in the cell division tree when an asymmetric division happens. These discretized states are sampled from all states such that the distance between two adjacent states on the cell state tree is the same. This distance, denoted as *step_size*, is a user-defined parameter used in the simulation and we will investigate the resulting cell populations with various *step_size* values.

Based on the cell state tree and the sampled states, we can simulate the cell divisions and the two types of information of each cell: the lineage barcode and the gene expression data. Each cell’s gene expression data is associated with a cell state sampled from the state tree and its lineage. The procedure of generating the gene expression data of a cell is illustrated in Fig. 2a. At each cell division event, we first probabilistically decide whether this is a symmetric division or asymmetric division. We use the parameter *p_a_* to control the probability for an asymmetric division to happen. For each new cell, we first generate a cell identity vector (CIV) based on the CIV of its parent, and its corresponding SIV. In the case of symmetric division, the two daughter cells inherit the same cell state as their parent’s, the CIV of each is produced by adding the CIV of its parent and a random walk distance on the edge which follows a Gaussian distribution N(0,1). In the case of asymmetric division, one daughter cell inherits the same cell state as the parent cells, whereas the other daughter cell will move to a future state on the cell state tree (see Methods and the pseudocode in Supplementary Note Sec. 5 for the exact process of calculating cells’ CIVs). To account for varying differentiation speeds, for every asymmetric division, we sample the number of states that the cell moves forward on the state tree from a discrete distribution with finite support from 1 to *max_step*, a tunable upper bound.

The cell identity vector (CIV) of each cell models the combined heterogeneities of lineage and cell type of the cell and it can be considered as the cell’s latent space representation (Fig. 2d). It is then used to generate the gene expression profile of the cell (Fig. 2a). The advantages of introducing low-dimensional vectors to represent cell identities over manipulating gene expression counts directly are: 1) The CIVs, as latent-space representations of single cells, are able to characterize the biological dependencies of single cells during cell divisions which are consistent with the true trajectories provided by the cell state tree and the cell lineage tree; 2) Performing low-dimensional random walks are informative enough to capture the variances in the transcriptomic data while at the same time making the simulation computationally efficient.

Apart from each cell’s CIV, we also generate a gene identity vector (GIV) for each gene following SymSim (where the GIVs was termed as gene effect vectors). The GIV of a gene represents how much the gene is affected by each factor in the CIV. The GIVs can then be considered as the weights on the CIVs, which together decide the expression pattern of that gene in that cell. The details of generating GIVs can be referred to in Methods.

Now with the CIV of a cell and the GIV of a gene, we can generate the true mRNA counts of this gene in this cell through a two-state kinetic model [21, 30], where the parameters of the kinetic model are calculated from the CIV and GIV (Methods). After generating the true mRNA counts of genes in cells, we follow the steps implemented in SymSim [21] to add technical noise to the data by simulating the major steps in the experimental steps to get realistic observed gene expression counts (Methods).

With the procedures introduced above, TedSim can generate scRNA-seq data with different compositions and patterns of cell types. For trajectory inference, we would like the present-day cells to cover all the sampled states in the cell state tree so that we can use the scRNA-seq data of the present-day cells to potentially better reconstruct the underlying cell state tree. However, in practice, some experiments may capture only the terminal cell states, or part of the intermediate states. In order to investigate how the cell type composition affects the performance of TI methods, parameters *p_a_*, the possibility of asymmetric division, and *step_size*, the interval used to sample intermediate states, are introduced in TedSim. These parameters allow TedSim to control the speed of differentiating into the terminal cell types and simulate datasets with different compositions of cell states. Given the same number of cell divisions, larger *p_a_* and larger *step_size* values result in more cells at the terminal states (Fig. 2f-h). The states sampled using *step_size*=0.25 and *step_size*=0.5 respectively are shown in Fig. 2f. In order to show the effect of these two parameters, we take all the cells at the leaves of the cell lineage tree, and classify them by the depth of their corresponding states in the cell state tree. In Fig. 2g we show the frequency of cells belonging to states at different depths in the cell state tree. With larger *p_a_* and *step_size*, most cells’ states are the terminal states with the largest depth (bottom right plot of Fig. 2g). In Fig. 2h-i, we show the UMAP visualization of the present-day cells using two combinations of *p_a_* and *step_size* values, where one leads to discrete populations and the other leads to continuous populations. Being able to simulate either continuous or discrete population of cells using the same model with different parameters is one major difference of TedSim from other simulators that cope with the two different patterns of gene expression data separately.

TedSim is able to generate synthetic scRNA-seq data with multiple cell types that mimics real datasets in multiple aspects. Existing simulators compare various statistical properties of simulated and real data, including the distribution of mean expression and the percentage of zeros in each cell [19,21,24]. However, these publications use only statistical properties within one cell type or averaged over all cell types of real datasets to perform the comparison. In addition to preserving the statistical properties of every cell type individually, we show that data simulated by TedSim also maintains the cross cell type relationships as in the real data.

To demonstrate this, we use TedSim to generate a scRNA-seq dataset close to the gene expression data of 10 selected cell types of zebrafish larvae at 5 days post-fertilization from [23] (Fig. 3a) (Supplementary Note Sec. 4.1). First, we learn a cell state tree from the real data, in order to preserve the cross-cell-type relationship. We calculate the pairwise Euclidean distances between the mean expressions of the cell types, and then run hierarchical clustering to get the cell state tree (Fig. 3b) (Methods). Second, in order to maintain the relative numbers of cells in each cell type, we assign probabilities for the choice of each branch at each branching point in the cell state tree. With the fitted state tree and branching possibilities, TedSim is able to generate data that has similar cell type composition and relative locations of clusters (Methods, Figure 3C-E). Fig. 3f shows that the proportion of cells in each cell type matches between real and simulated data. By tuning the parameters across multiple stages of the simulation, we can match the statistical properties of the simulated data to a given dataset (Methods). Fig. 3g-h show respectively the comparison between simulated and real data in terms of mean gene expression and percentage of zeros per cell in each cell grouped by cell type, and the gene-wise comparisons can be found in Supplementary Fig. 1.

**Figure 3.**
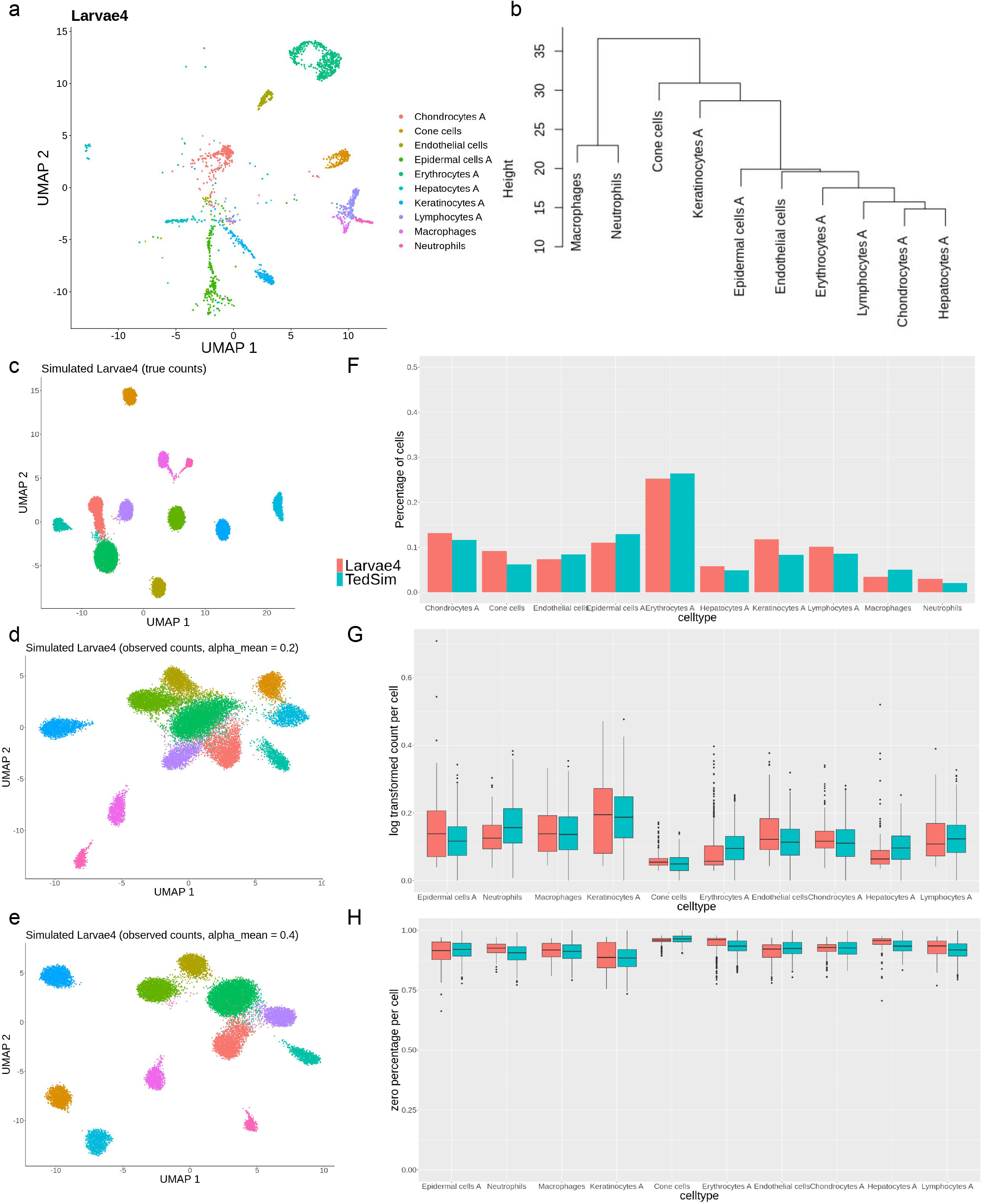
Fitting TedSim to a real dataset. (a) UMAP visualization of 5-dpf zebrafish larvae cells from (Spanjaard and Hu et al. 2018). 9393 cells of 10 cell types are sampled from the original dataset. (b) Cell state tree estimated from the real dataset. Hierarchical clustering is applied to the pairwise distances of the average expressions of the 10 cell types. (c - e) UMAP visualization of simulated zebrafish larvae cells. 20 simulations each with 1024 cells are generated with tuned parameters to fit the dataset to the referred real dataset. (c) The true counts. (d and e) The observed counts with varying capture efficiency. (f) Comparison of cell type compositions between simulated data and real data. (g-h) Statistical comparison between cell types of simulated data and the real data. Both metrics are calculated based on observed counts with capture efficiency *alpha_mean* = 0.1. (g) Average expression per cell. (h) Average zero percentages per cell.

### Simulating heritable lineage barcodes during cell divisions

TedSim simulates the cell lineage tree, which is a binary tree that models the cell division events (Fig. 4a). Starting from the root cell, we simulate the accumulation of CRISPR/Cas9 induced scars along the paths from the root to all the leaf cells. In practice, the barcodes with accumulated scars in present-day cells can be obtained using scRNA-seq and used to reconstruct the cell lineage tree (Fig. 4a).

**Figure 4.**
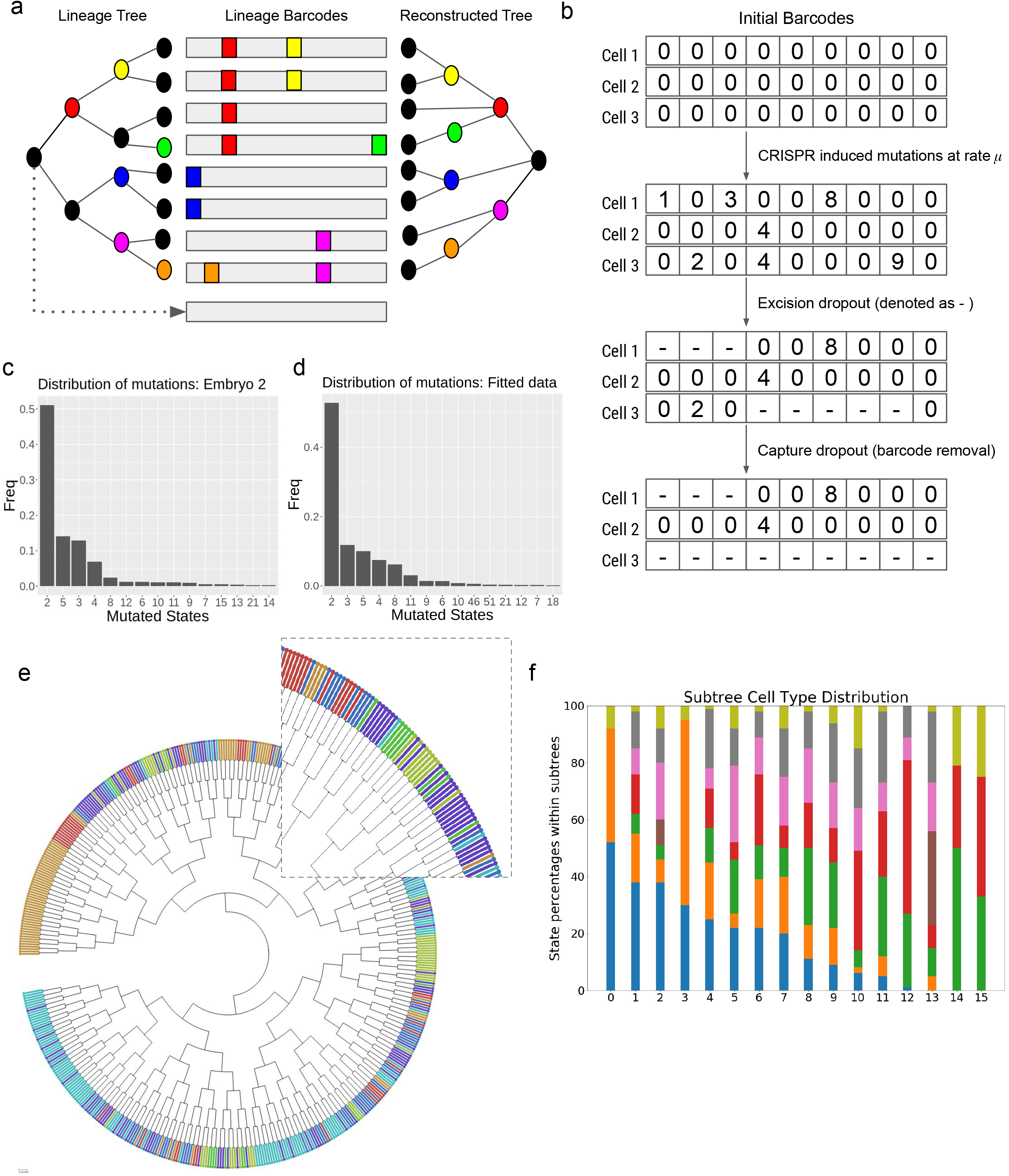
Simulation of lineage barcodes and inconsistency between cell lineage and cell state. (a) Reconstructing cell lineage using simulated lineage barcodes. During the cell division process, mutation events can randomly happen (as colored on the left) and cause insertion/deletion on the lineage barcode (middle), a colored box represents an insertion/deletion and is termed a mutated state (non-zero characters in the character vectors). Given the mutated barcodes of the present-day cells (leaf nodes of the lineage tree), tree reconstruction algorithms can be applied to infer the lineage tree (right). (b) Simulating dropouts of lineage barcodes. When two or more mutations happen at one division, the excision dropout will randomly drop characters between two mutated sites; after library preparation steps, the capture dropout will completely drop the barcode if the attached gene is not captured. (c-d) Distribution of mutated states in the lineage barcodes. (c) Mutated state distribution (only top 15 most frequent mutations are shown) observed in lineage barcodes of the embryo 2 from M. Chan et al. [14], and the frequencies are used to generate synthetic mutations in TedSim. (d) Mutated state distribution (top 15 frequent mutations are shown) in simulated lineage barcodes by TedSim. Mutation rate *μ* = 0.1. No dropouts are introduced. (e) Cell type visualization onto the lineage tree shows inconsistency between states and lineage barcodes. Different Colors represent different cell types. (f) Relative cell type distribution descended from ancestor cells at the same depth (the fifth generation from the root) in (e).

The lineage barcodes of a cell can be represented by a character vector of which each character represents a target where a CRISPR-induced mutation can possibly happen [14, 22] (Fig. 4b). The root cell in the cell division tree has its default barcode with all states unmutated (an unmutated state is denoted as “0”) . When it divides, mutations happen at a mutation rate *μ* that randomly select certain characters and change them to some mutated states (denoted as non-“0” integers) in the daughter cells, and the mutations will be carried on to further descendants (Fig. 4b).

Dropouts in the lineage barcodes is one of the main challenges towards reconstructing the cell division tree using computational algorithms. Following Salvador-Martínez *et al* [22], TedSim models two kinds of dropouts widely occurring for in-vivo experiments (Fig. 4b): ‘capture dropout’ that refers to experimental dropouts due to low capture efficiency, and ‘collapse dropout’ or ‘excision dropout’ typically means inter-target deletions when two or more sites are cut at the same time [31]. Both dropouts are denoted as ‘-’ in the character vectors. According to certain experimental protocols where the barcode is inserted to a specific endogenous site (a real gene or an artificial gene) in the genome [32], we associate the capture dropout of the lineage barcode in each cell with the expression level of the gene in the cell. If the gene is not captured in the simulated observed counts, the capture dropout to the lineage barcode is applied. Simulating gene expression profiles together with lineage barcodes allows TedSim to generate more realistic dropout events compared to randomly removing barcodes. A number of mutation events can not be observed due to dropouts, which limits the accuracy of cell division tree reconstruction. In TedSim, users can choose whether or not to include dropout in the simulation using a boolean parameter *p_d_*.

Another phenomenon observed in real data is that the mutations do not occur at every position of the barcode with equal probabilities, but rather show significant biases towards certain mutations and the frequencies of the mutations follow an exponential curve [14, 22] (Fig. 4c). In TedSim we simulate this bias by sampling the mutation events from such distribution obtained from an experimental dataset in [14]. Therefore, we are able to generate simulated lineage barcodes that have a similar pattern of mutation distribution as the real dataset (Fig. 4d). In the TedSim implementation, we also keep the alternative of uniformly distributed mutated states which can help to test tree reconstruction algorithms under ideal circumstances where mutations occur uniformly at different positions of the barcode. Later in this paper we will show that the differences between the two distributions of mutations can influence the performances of tree reconstruction algorithms.

Given the simulated lineage barcodes, reconstructing the lineage tree is an NP-hard problem [33, 34]. The state-of-the-art tree reconstruction algorithms tend to have no guarantee of how close the solution is to the real lineage tree. By comparing the ground truth and the reconstructed tree, TedSim is able to benchmark the performances of different methods. In comparison to [22], TedSim has the following new features in terms of simulating lineage barcodes: 1) TedSim simulates lineage barcodes simultaneously with gene expression data (CIVs) while traversing the cell lineage tree; 2) The simulated lineage barcodes can be connected to a gene where the capture dropout can happen based on the observed count of the gene. These features make it more feasible to use the combined profile generated by TedSim to benchmark multimodal methods.

### TedSim generates the inconsistency between transcriptome similarity and barcode similarity observed in real data

One important question in developmental biology is how cell types are related to cell lineages.With the development of CRISPR/Cas9 lineage barcoding alongside scRNA sequencing, we are able to uncover the emergence of complex multicellular organisms from single totipotent cells. Such molecular recording techniques have been able to provide insights into how the transcriptomic data and the cell lineage are related by examining the cell type composition within subtrees at different levels of the cell lineage. Such lineage recorders [14, 23] have shown that the relative cell distribution can be mixed with cell types while at the same time biased towards some cell types, which may be caused by positional biases [35, 36]. Basically, existing data show that there is a considerable number of cells of the same type which are located in different subtrees of the reconstructed cell division tree, and in the same subtree there can be multiple cell types [13,14,23,37]. We call this the inconsistency between transcriptome similarity and lineage barcode similarity. Various scenarios that cause this inconsistency are discussed in [10].

The model we use in TedSim combining an underlying cell state tree with asymmetric cell divisions can generate such inconsistency between transcriptome similarity and barcode similarity of cells. TedSim assumes, in asymmetric divisions, that different paths of cell fates are chosen randomly with certain possibilities, and since the state shifts can only go from the root to the terminal states, the random choice of early-stage cells can result in significant biases towards some cell fates. Therefore, TedSim can model the realistic cell type composition within subtrees of the lineage tree (Fig. 4e-f). Chan *et al* presented the cell type composition under various progenitors (Fig. 6a in their paper) [14] in a similar form to Fig. 4f, from which we see that the simulated dataset shows similar heterogeneity in cell types in each subtree/progenitor, including a mix of multiple cell types as well as biases towards some cell types.

The connections between lineage and transcriptomic data can also be dynamic and transient [28]. By default in TedSim, we assume that the state of the cell has a bigger impact on the gene expression than the lineage of the cell, and we are able to achieve different ratios by tuning the weight of the SIVs in the CIVs (Methods). With future research uncovering how the correlation between cell lineage and cell state changes over time, we will be able to easily generalize the current model to automatically change the weight of SIVs in CIVs based on the developmental stages the cell is on.

### Benchmarking trajectory inference methods

ScRNA-seq data provides transcriptome information of every single cell, but as a cell can be sequenced only once and in most of the experiments all cells are sequenced at the same time, scRNA-seq data does not provide data of the cells in the past for researchers to study developmental processes. Trajectory inference (TI) methods were developed to infer the cell trajectories using only data from present-day cells, assuming that there are cells in the dataset which represent cell states at various stages of the developmental process.

Various TI methods have been benchmarked in multiple publications using simulated data [1, 21], but the data simulation procedures used in these publications align with the assumption that the data covers the entirety of the continuous manifolds that the transcriptomic data lives in. On the contrary, TedSim generates scRNA-seq data through realistic cell division events that do not guarantee that the states of the cells simulated cover all the sampled states on the cell state tree, therefore, the simulated data can reflect the realistic challenges for the TI methods. In particular, in previous sections we showed that the cell type composition in the present-day cells can be different depending on various factors (Figs. 2F-I), and how balanced the cell state composition is in the data can be important for the performance of TI methods. Therefore, instead of comparing different TI methods, we will focus on investigating the performances of the top TI algorithms on datasets with different compositions of cell states, controlled by TedSim parameters.

When applying a TI method to data generated by TedSim, the input to the TI method is the scRNA-seq data from TedSim, and the cell state tree used in TedSim serves as ground truth to evaluate the TI method (Fig. 5a). We apply two state-of-the-art TI methods, Slingshot [2] and PAGA-Tree [3], as benchmarked by [1]. Fig. 5b shows the inferred trajectory by Slingshot on one of our datasets generated by TedSim with well tuned parameters, which indicates that with proper settings to cover most of the cell states in the data, the TI methods are able to infer the correct state tree shown in Fig. 5a. We then focus on testing TI methods under various compositions of cell types in the present-day cells, which can be controlled by parameters *p_a_* and *step_size*. We vary parameters *p_a_* and *step_size* when generating and applying PAGA-Tree on the resulting simulated scRNA-seq datasets. Fig. 5c shows the UMAP visualization of the simulated data and inferred trajectories. Consistent with Fig. 2f-i, high *p_a_* and *step_size* result in more discrete populations of cells, mainly belonging to the terminal states in the cell state tree. Lower *p_a_* and *step_size* values, on the other hand, lead to more cells at non-terminal cell states. TI methods work better when the present-day cells cover as many cell states (both terminal and non-terminal) as possible. For example, PAGA-Tree infers the correct trajectory when pa=0.6 and step_size=0.25, while when pa=1 and step_size=1, it is very difficult for the TI method to find the correct trajectory. The results of Slingshot on the same datasets are shown in Supplementary Fig. 2.

**Figure 5.**
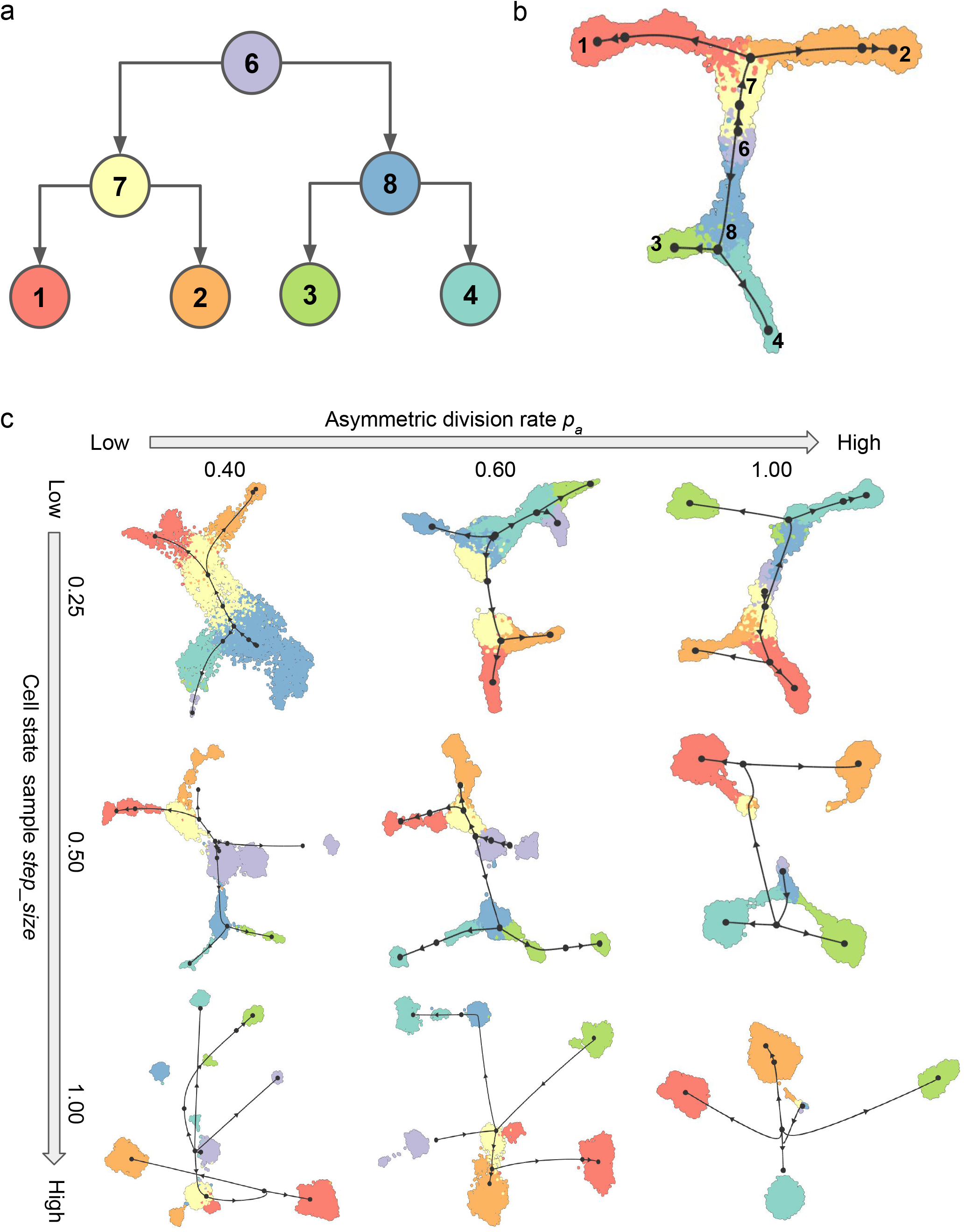
Running trajectory inference methods on TedSim datasets. (a) The structure of the cell state tree used for the simulation used in this figure. Number of intermediate states on each edge varies with *step_size*. (b) UMAP visualization of inferred trajectories by Slingshot, on a dataset with 8192 cells and 500 genes. *p_a_* = 0.85 and *step_size =* 0.25; these parameters are selected to obtain continuous populations of cells. (C) UMAP visualization of inferred trajectories by PAGA-Tree when varying *p_a_* and *step_size*.

**Figure 6.**
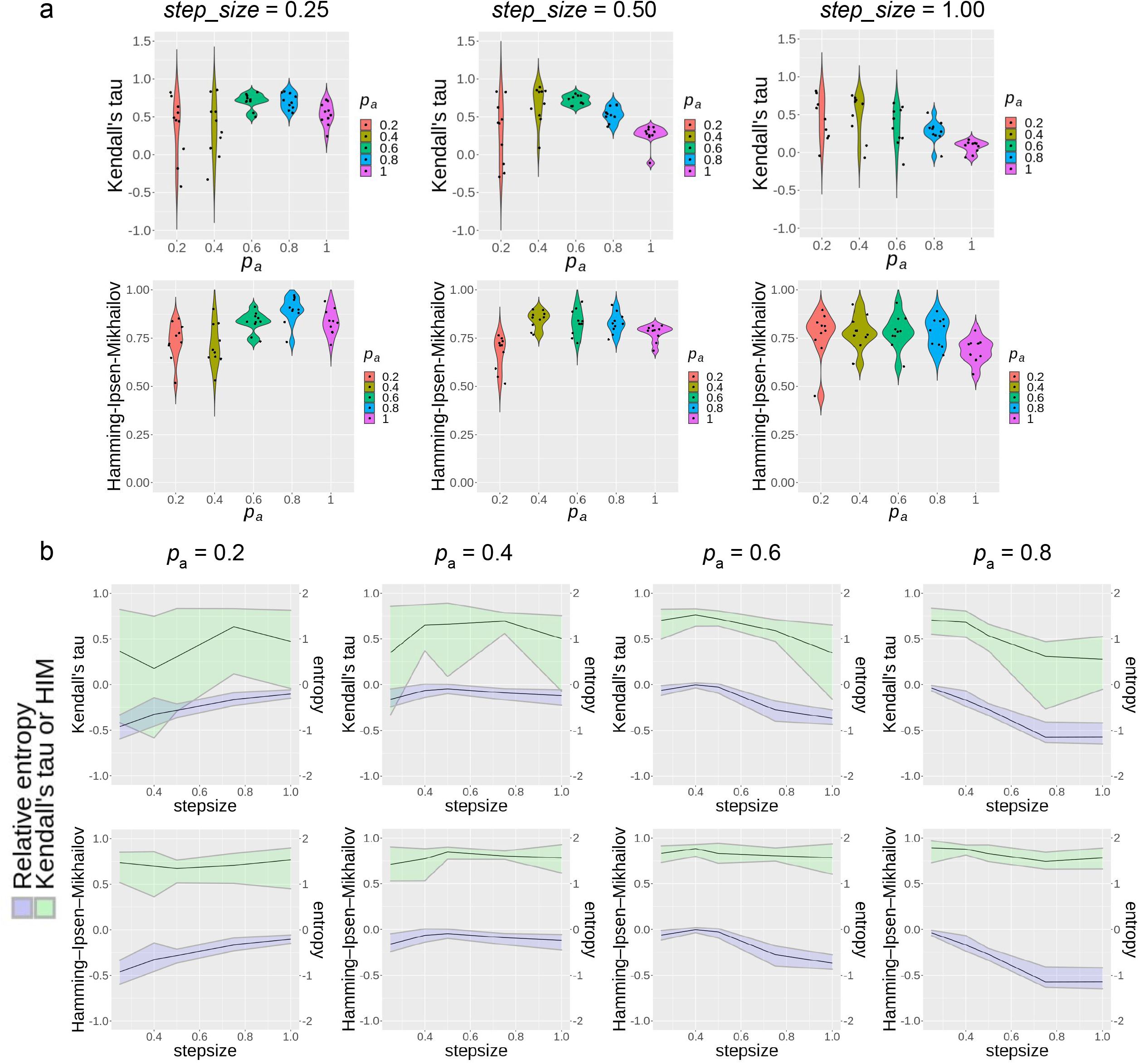
Statistical results of trajectory inference methods. (a-b) Evaluating PAGA-Tree results on varying parameters *p_a_* and *step_size*. 10 datasets are simulated for each parameter configuration to generate the violin plot. (a) Pseudotime inference performances with the same *step_size* and varying *p_a_* . The ground truth for pseudotime rank is obtained from the depth of the cells’ states on the state tree, and the HIM is calculated between the state tree topology and the milestone network (Saelens et al. 2019) inferred by the TI method. (b) Pseudotime inference performances of PAGA-Tree and relative entropy of the dataset with varying *step_size* parameters. The black curve represents the average score of 10 runs and the upper and lower bound of the ribbon represents maximum and minimum score.

To quantitatively measure how balanced the cell type composition is in a dataset, we propose *relative entropy*. The relative entropy first calculates the entropy of the discrete probability distribution derived from the frequencies of each cell type, then normalizes this entropy against the length of the distribution (ie. the total number of cell types) to remove the bias of probability distribution length on entropy (Methods, Supplementary Note Sec. 2). Higher relative entropy corresponds to more balanced populations. We also quantitatively evaluate the performance of the TI methods under different *p_a_* and *step_size* values. We calculate Kendall’s *τ* [38] correlation between the inferred pseudotime and true pseudotime and the Hamming–Ipsen–Mikhailov distance (HIM) [39] for the difference between the topology of inferred and true trajectories as used in [1]. We plot the relative entropy of the datasets along with the Kendall’s *τ* and HIM score of trajectories inferred by PAGA-Tree in Fig. 6b. We observe that Kendall’s *τ* changes in consistent with the trend of the relative entropy, where for smaller *p_a_* such as *p_a_* = 0.2, the relative entropy increases with *step_size* since it requires larger *step_size* in order to capture the whole developmental trajectory; on the contrary, when *p_a_* gets closer to 1 and cells get to the terminal states fast, the relative entropy declines with *step_size*. The HIM score does not change much along with the change of parameters, which indicates that with a given starting cell of the trajectory, the continuity of the dataset does not affect the inference of the topology of the state tree as much. The quantitative results of Slingshot on the same datasets can be referred to in Supplementary Fig. 3.

From Supplementary Fig. 3, which includes results from both Slingshot and PAGA-Tree, one can compare the performance of these two methods in different scenarios. Overall, Kendall’s *τ* values of the two methods are comparable, but PAGA-Tree outperforms Slingshot in recovering the structure of the state tree, measured by the HIM score. The results indicate that TedSim is able to model the hidden factors that affect how the pattern of scRNA-seq data changes, and is able to evaluate the trajectory inference methods from multiple perspectives of how accurate it assigns pseudotime to single cells, as well as how well it preserves the underlined state tree under different speeds of differentiation.

These results suggest that the TI methods are likely to find the correct underlying trajectory which represents cell state transition if the dataset covers most states on the state tree and forms continuous populations. If the dataset contains only terminal cell states, the TI methods can not find the correct trajectory. Therefore, when applying TI methods, it is important to be aware that the composition of cell types in the dataset is a key factor to determine the reliability of the results.

TedSim has the potential to suggest the time to harvest cells for the most comprehensive cell type compositions. For example, given a hypothesized asymmetric cell division rate and cell state tree, running TedSim allows for the estimation of the number of divisions needed to obtain cells with comprehensive cell type compositions.

### Benchmarking hybrid methods of various inference tasks

With the development of technologies that jointly profile lineage barcodes and scRNA-seq data, hybrid computational methods are emerging to integrate these data for various purposes. Three hybrid methods, lineageOT [18], TreeVAE [40] and LinTIMaT [17] are tested in this paper, where the first two are benchmarked in this section.

Given time course scRNA-seq data, researchers try to infer the relationships (the ancestor-descendent correspondence) between cells at different timestamps [8]. Forrow and Schiebinger proposed LineageOT [18], which aims to incorporate the cell lineage tree reconstructed from the lineage barcodes to improve the accuracy of ancestor-descendant inference when the lineage barcode data is available. Here we use the data simulated by TedSim to benchmark this method (Methods). As performed in [18], we compare their proposed method LineageOT with EntropicOT [8] which uses only gene expression data. From the results, we can see that by using good lineage information (LineageOT, true tree), the normalized ancestor error can be largely improved. By tuning the parameters which affect the continuity of populations and cell type compositions *p_a_* and *step_size*, we can also see how the quality of the gene expression data influences the ancestor inference of all three methods. As shown in Fig. 7a, all methods perform worse when *p_a_* and *step_size* are increased, considering the fact that larger parameters result in more discrete populations. In this case, the cells at two different generations are more likely to have the same cell state, making it harder to infer the ancestor-descendant relationships. While LineageOT with true lineage tree is still able to maintain decent ancestor error, LineageOT with fitted tree falls behind even compared with EntropicOT. How to better reconstruct the lineage tree from the barcodes will be the key to infer the developmental trajectory more accurately.

**Figure 7.**
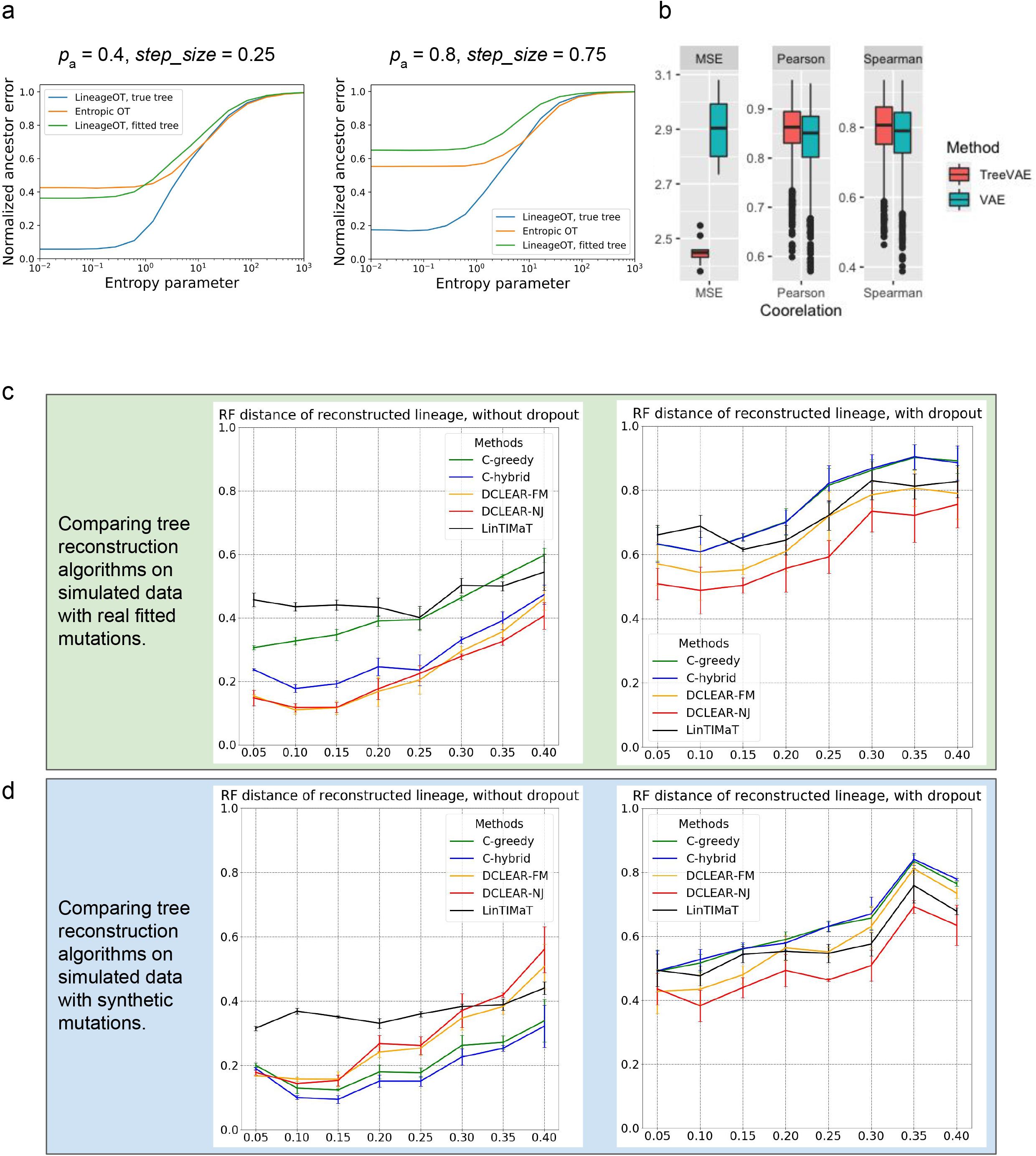
(a) Benchmarking ancestor-descendant inference accuracy of LineageOT. The normalized ancestor error is calculated as the marginal of the mean squared optimal transport distance between inferred ancestors and ground truth [18]. (b) Benchmarking ancestral gene expression inference accuracy of TreeVAE. The correlation and distance metrics are calculated between the inferred gene expression of ancestor cells and the true counts of the same cells given by TedSim. (c-d) Benchmarking tree reconstruction methods on synthetic lineage barcodes using Robinson-Foulds distance. Tree reconstruction accuracy of 5 selected methods on simulated lineage barcodes. The mutation states are sampled from a fitted distribution of real mutations of embryo2 from M. Chan et al. (c) or a uniform distribution of synthetic mutations (d) Methods except LinTIMaT utilize only the lineage barcodes while LinTIMaT also uses gene expressions. Parameters used to simulate the datasets can be found in Supplementary Note Sec. 4.3. For each configuration (mutation rate and dropout), 10 simulations are run and used to test the methods.

Another computational problem using both the scRNA-seq and the lineage tracing barcode data is to reconstruct the gene-expression profiles of ancestral cells on the cell division tree. Ouardini *et al* developed TreeVAE, which utilizes paired single-cell lineage tracing and transcriptomics data [40]. Here, we test the accuracy of the reconstructed gene expressions of ancestor cells using the known ancestral data in TedSim. The method is compared with a VAE (variational auto-encoder) -based baseline method, which first runs on leaf nodes, then calculates the gene expressions of an internal node by averaging the latent space for leaves below the node and decode it using the trained decoder [40]. With the ground truth of the ancestor cells’ gene expression, we evaluate the inferred expression of all ancestor cells by calculating the Pearson’s correlation, Spearman’s correlation and MSE (mean squared error) scores (Fig. 7B). From the results, we can see that TreeVAE outperforms the baseline method in all three metrics. Besides, we also compare the inferred expressions for individual genes using either the true lineage or reconstructed lineage from the simulated lineage barcodes. From the results, we can see that the correlation scores of TreeVAE still consistently outperforms the baseline method no matter which tree is used (Supplementary Fig. 4).

### Benchmarking lineage reconstruction algorithms

TI methods applied to single-cell gene expression data can, to a certain extent, reconstruct the change of cell states during developmental processes. The lineage tracing barcodes, on the other hand, are expected to infer the cell lineage tree, i.e., the cell division history and the clonal relationships between cells. A few computational methods have been developed to reconstruct the cell lineage tree from the lineage barcodes of the present-day cells [16, 22]. However, it is very challenging to obtain a cell lineage tree with high accuracy due to various reasons including the large number of cells, the limitation of target sites and missing data in the barcodes [22]. To improve the accuracy of reconstructed trees, hybrid methods that use both the scRNA-seq and lineage barcode data are emerging, with LinTIMaT [17] as a representative. Since TedSim generates both scRNA-seq and the lineage barcode data simultaneously, it can be used to benchmark not only tree reconstruction methods that use only lineage barcode data, but also those that use both gene expression and barcode data. In particular, we would like to find out whether the performances of lineage reconstruction are improved by integrating gene expressions with the lineage barcodes. We compare the performance of LinTIMaT [17] against DCLEAR [41] and Cassiopeia [16], which are the best performing algorithms in the recent Allen Institute Cell Lineage Reconstruction DREAM Challenge [41] which use only the CRISPR/Cas9 induced lineage barcodes.

We set up the simulation of TedSim as follows: for each cell, the lineage barcode contains 32 target sites, and each target site, if mutated, can be sampled from 100 possible mutated states. Simulations are run in varied mutation rate *μ* and dropout rate (on/off), and for each configuration we repeat the simulation 10 times. The parameters for simulating gene expression values are included in the Supplementary Note Sec. 4.2. To evaluate performances of the methods, we use two measures: (1) Robinson-Foulds (RF) distance [42] that quantifies the symmetric difference in splits between the reconstructed tree and the ground truth, which reflects the accuracy of the overall topology of the tree; (2) Triplets correct rate that calculates the percentage of correctly located triplets of cells on the cell lineage, which reflects the accuracy of inferring local lineages of single cells.

From the benchmarking results (Fig. 7c), we observe that while RF-distance metric for all methods shows consistent dependency on the mutation rate *μ*, and the trend is consistent with that is shown in [22] although Salvador-Martinez *et al* used different lineage tree reconstruction methods. The triplet correct rate results are much less affected by *μ* (Supplementary Fig. 5). Cassiopeia-hybrid has the best triplet correct rates overall, especially outperforms all other methods by a large margin under the condition of high mutation rate without dropout, possibly due to the advantage of using an ILP solver. With dropout, Cassiopeia-greedy outperforms Cassiopeia-hybrid slightly, but both are still better than the two DCLEAR modes and LinTIMaT. The results of RF-distance show similar trends for all selected methods, and we find that the best mutation rate is around 0.1-0.15 for both conditions: with and without dropouts, and excessively high mutation rate does not result in higher tree reconstruction accuracy as people may intuitively think so. In Fig. 7d, we show the same comparison of tree reconstruction methods using simulated data where the mutations occur uniformly on the barcode. Comparing Fig. 7c and Fig. 7d, we see that the tree reconstruction methods generally perform better with the data where mutations are uniform, suggesting that the bias in mutation distribution on the barcode should be modeled when developing tree reconstruction methods in the future.

Interestingly, we see that LinTIMaT does not perform well for lower mutation rates but its performances decline much slower with the increase of mutation rate. The reasons that LinTIMaT does not overperform other methods, as we interpret, are two folds: (1) the initial tree obtained by LinTIMaT before it is refined using gene expression data is of low quality, which can be due to bad initialization, and this affects the accuracy of the final output tree; (2) LinTIMaT relies on the assumption that cells with similar gene expression profiles should be located closely in the cell lineage tree, which is not always true. On the other hand, LinTIMaT’s performance does not deteriorate as much as other methods when the mutation rate increases because the incorporation of gene expressions helps to maintain relatively good results when the barcodes are not very informative.

## Discussion

We presented TedSim, a simulator which generates both scRNA-seq data and lineage barcodes simultaneously through simulating the cell division events. Compared to existing simulators of scRNA seq and CRISPR lineage recorders, TedSim has the following novel features: (i) TedSim models the underlying temporal dynamics of the development of cells by simulating the processes of cell division and differentiation to get both gene expressions and lineage barcode data simultaneously. (ii) TedSim is able to simulate both discrete and continuous gene expressions of single cells under the same framework by adjusting the cell differentiation speed and the continuity of cell states. (iii) Given simultaneously simulated gene expression and lineage barcode data, TedSim can benchmark hybrid methods which use both types of data, including cell lineage reconstruction methods which incorporate gene expression data (LinTIMaT) and trajectory inference methods that use both time-series gene expression data and cell lineage tree (LineageOT). (iv) The simulated data shows realistic inconsistency between transcriptome similarity and lineage barcode similarity of cells, which cause the heterogeneity of cell types in a lineage subtree, and meanwhile the subtree can have a dominant cell type.

By applying state-of-the-art trajectory inference methods to scRNA-seq data simulated by TedSim, we have gained insights towards experimental design for studying cell trajectories: it is important to maintain cells from ancestral states in the cell state tree and the cell type composition in a dataset is crucial for the TI methods to return meaningful inference. One may consider sequencing cells at multiple time points in order to capture early cell states. Overall, we found that the TI methods, although using scRNA-seq data of present-day cells, can recover the underlying state tree that guide the cell division and differentiation events reasonably well, as long as the dataset has a good coverage of both terminal and non-terminal cell states.

As more datasets which profile both the CRISPR/Cas9 induced scars and single cell gene expression data become available, hybrid algorithms that make use of both types of data to learn cell dynamics are needed. Current hybrid methods (for example, LinTIMaT), as we benchmarked with TedSim, have not provided satisfying results. Future methods may benefit from better modeling the relationship between the two types of data, possibly using the asymmetric cell division model we propose in TedSim.

## Materials and Methods

### Simulating State Identity Vectors with the Brownian motion model

In TedSim, we consider the scenario where cells differentiate following a tree structure. The cell state tree represents the structure of this differentiation (Fig. 1). Users can define the desired structure of the tree by inputting the tree in Newick format in a text file. State Identity Vectors are introduced to determine heterogeneity (by diff-SIVs) as well as homogeneity (by nondiff-SIVs) between cell states. The differences between cell states are realized through diff-SIVs, and the user is able to define the number of diff-SIVs and nondiff-SIVs. Nondiff-SIVs, whose behavior are consistent across cell states, are sampled from a Gaussian distribution with a given mean (default is 1) and variance (default is 0.5). The changes of diff-SIVs along the paths of the cell state tree are modeled by Brownian motion along the tree from root to leaves, where one starts with a given value at the root (default is 1), and at each time point t, y(t) is calculated as y(t)=y(t−Δt)+N(0,Δt), where N(·) represents a Gaussian function, and Δt is the step size of the random walk. The values at branching nodes of the tree are shared by all branches connecting this node. The Brownian motion model is able to capture the structure of the tree where the expected distance between diff-SIVs of two states is proportional to the total edge length it takes to walk from one state to another (Supplementary Note Sec. 3). We consider the values sampled by the Brownian motion model as one feature of the identity of cell states. A number of features that define the state identity, termed as State Identity Vector, are generated by running the Brownian motion on the cell state tree repeatedly. After running Brownian motion on the state tree to get the SIVs on the state tree (Fig. 2c), we sample the SIVs based on a constant parameter *step_size* and keep only the sampled states as the possible cell states of the cells. The states of the cells are chosen from the sampled states when iterating on the cell division tree through symmetric and asymmetric division mechanisms.

### Simulating Cell Identity Vectors with the Brownian motion model

When generating cells through cell divisions, TedSim models the identities of single cells using Cell Identity Vectors (CIVs), which adds up two modes of variations from both the cell state tree and the lineage tree.

Given the cell state tree and the cell division tree, the states of all the cells can be determined using a two-mode division strategy: symmetric division, meaning no state changes during division; asymmetric division, where one of the daughter cells will be shifting from the current state of the parent to the future sampled state on the state tree. Starting from the root of the cell division tree, we first initialize its state to be the root on the state tree. Recursively, we determine the states of the children based on the state of their parent cell, with either symmetric division or asymmetric division. We denote the possibility of a cell to divide asymmetrically as *p_a_*. The asymmetric division rate *p_a_* influences how fast the cells can get to the terminal cell states (Supplementary Note Sec. 1). Once the asymmetric division occurs, one of the daughter cells will trigger a state shift. The step size of one state shift is sampled from a discrete distribution with a finite support from 1 to *max_step*, a tunable upper bound. When there is a split of branches on the state tree, the state shift will randomly select an edge to follow, directed by branching possibilities. The branching possibilities determine the balance of the cell types under two subtrees. In order to fit the cell type composition from a real dataset, the branching possibilities are calculated by weighing the total number of cells under the subtrees (Supplementary Note Sec. 5). The combination of symmetric divisions and asymmetric divisions, which allow fast differentiation, models the biological processes which lead to multiple cell types including un-differentiated cell types in a dataset with only present-day cells.

With SIVs simulated by the Brownian motion and cell states determined by the two-mode division strategy, TedSim simulates CIVs using the Brownian motion model. To be more specific, Given the CIV of the parent cells y_p_(s), where s is the state of the cell, the CIVs of daughter cells can be calculated using a function as: y_c_(d(s)) = y_p_(s)-SIV(s) + SIV(d(s)) + N(0, 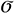^2^) = y_p_(s) + N(SIV(d(s)) - SIV(s), 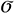^2^), where d(.) represents the two-model division function to decide the state of the cell, SIV(.) is the SIV value of the state, N(0, 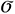^2^) is the Brownian motion term that represents the random walk distance from the parent to the daughter cell on the cell division tree where 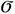 is kept constant (default as 1) since the edges of the lineage tree are considered the same. The difference of generating CIVs from generating SIVs that both utilize a Brownian motion model is that the random walk samples from a Gaussian function whose mean is determined by the state changes of the daughter cell from the parent cell and standard deviation reflects the lineage distance between the parent and the daughter cells.

### Simulating true mRNA counts and observed gene expression

Following SymSim [21], we consider that each gene has an identity, denoted by a *gene identity vector* (GIV, referred to as “gene effect” vector in SymSim). The GIVs are first drawn independently from a normal distribution, and then dropped to zero with probability η, making sure that each gene is only affected by a small subset of positions in the CIV. The CIV of a cell and the GIV of a gene together determine the gene expression in the cell. Similar to SymSim, TedSim simulates the true mRNA count of a gene in a cell via the two-state kinetic model [30, 43], and the parameters of the kinetic model (*k*_on_ and *k*_off_ and *s*) were calculated from the CIV and the GIV. For a gene in a cell, it can be either *on* or *off* and the parameter *k*_on_ represents the rate the gene being switched on and *k*_off_ the rate gene being switched off. When a gene is in the *on* state, its transcript is synthesized with rate *s*, and the generated transcript then degrades with rate *d*. When *d* is fixed, the distribution of the mRNA counts can be modeled with a Beta-Poisson distribution [43], and we can sample the mRNA count of the given gene in the given cell from the distribution with calculated *k*_on_ and *k*_off_ [21].

Single-cell gene expression data obtained with scRNA-seq technologies are confounded by technical noise. The technical noise is introduced during sample processing steps such as mRNA capture, reverse transcription, PCR amplification, RNA fragmentation, and sequencing. TedSim simulates the key technical steps to generate observed gene expression levels. Following SymSim [21], TedSim simulates two categories of library preparation protocols, one does not use UMIs (unique molecular identifiers) [44] and sequences full-length mRNAs (using procedures in Smart-seq2 [45] as template), and the other uses UMIs and sequences only the 3′ end of the mRNA (using the Chromium chemistry by 10x Genomics as template).

### Fitting simulated gene expression data to real datasets

We demonstrate how TedSim can generate realistic gene expression data that shares cell-type specific properties with real datasets where there are multiple cell types. We use a zebrafish larvae dataset at 5 days post fertilization as the reference dataset [23]. We removed undifferentiated cell types and selected 10 cell types with 9393 cells as terminal cell states (Supplementary Note Sec. 4.1). We then generate simulated data with TedSim that resemble the properties of the real data in terms of variation between cell types and gene expression distribution in each cell type, using the following procedure.

First, we used Ward’s agglomerative clustering method to obtain the cell state tree (implemented in “The R Stats Package”). The edge lengths of the state tree are rounded up to the nearest integer. With the inferred state tree, in order to maintain the relative size of each cell type (that is, the proportion of cells in each cell type), at each branching node in the cell state tree, we assign probabilities of choosing each branch according to the total number of cells in the respective subtree (Supplementary Note Sec. 5). We ran the TedSim simulation 20 times, each with a single root and having 1024 leaf cells, using the same cell state tree and branching possibilities. We then combine the cells from the 20 runs and obtain the final dataset. The asymmetric division rate *p_a_* and *step_size* used in the simulations are tuned to achieve discrete populations of the cell types so that the numbers of cells of terminal cell types are maximized (*p_a_*=1, *step_size*=1). Besides, we calculate the mean gene expression per cell type, and fit the *scale_s* parameter, which was also used in SymSim, to adjust for the difference in cell size of different cell types such that the total UMI counts in each cell when stratified by cell types match the same pattern of that in real data (Fig. 3g). Similarly, the percentage of zeros in each cell, affected by both the cell size and the capture efficiency, also achieves similar statistical properties to the real dataset in each cell type (Fig. 3h). The detailed information about the parameters used for the simulations can be found in (Supplementary Note Sec. 4.1).

### Simulating CRISPR/Cas9 induced lineage barcodes simultaneously

As we generate CIVs of the cells along the cell division tree, the lineage barcodes are also generated simultaneously. The lineage barcode of a cell is represented as a vector of target sites where different mutation events are represented by a unique character (Fig. 4b). The lineage barcodes of the daughter cells are derived based on the parent’s barcode through possible mutation events. Following [14], we denote the unmutated state as ‘0’ and mutated states as non-zero integers in the character vectors. We keep the mutation rate *μ* constant across all target sites and all cell divisions, then the probability of a particular mutation happening follows a geometric distribution. We assume that the mutation is irreversible so that a mutated target will not go back to the unmutated state.

The possible mutation states are either synthetically generated or fitted from a real dataset, and the number of mutated states is controlled by the user. By default, if we do not learn the mutated states from a real dataset, the mutated states will be uniformly distributed, achieving maximum entropy for a fixed number of mutated states. On the other hand, given the character matrix (cell by barcode) of a real dataset, we fit the mutation events by first calculating the frequencies of each occurred mutation, and then taking the top *N*_ms_ most frequent mutations (*N*_ms_ is a user defined parameter). We increase the frequency of the last mutation state such that the total frequency sums up to 1.

TedSim incorporates two kinds of dropouts: excision dropout and capture dropout. To generate excision dropout, if two mutations happen during one division, all the targets in between will drop to ‘-’, which is the missing state. If three or more mutations occur, we will randomly select two of the mutated states and then remove the targets in between. The capture dropout is introduced due to the low capture efficiency during sequencing. We first calculate the averaged observed expression level of all genes, and bind the lineage barcodes to one of the highly expressed genes. The barcodes of cells with zero count of that gene will then be removed, which means that all targets will drop to ‘-’. This process can be considered stochastic since the capturing step in the library preparation step captures the molecules with a stochastically generated efficiency parameter. The pseudocode for generating lineage barcodes simultaneously with CIVs can be found in (Supplementary Note Sec. 5).

### Evaluation of trajectory inference methods

The cell state tree used in TedSim is the ground truth cell trajectory for trajectory inference methods. When using TedSim to benchmark TI methods, we investigate the performance of TI methods with cell populations of different differentiation speeds by tuning the parameters *p_a_* and *step_size*. For each combination of parameters, we simulate 8192 cells with the same bifurcating lineage and run the simulation 10 times to calculate the average performance. Values of other parameters used to generate the datasets can be found in (Supplementary Note Sec. 4.2)

With a simulated dataset, we first apply PCA and select the first 50 PCs to input to UMAP for visualization in 2-d, and set min_dist = 0.3 to maintain local structures.

We use the R packages dynwrap (https://github.com/dynverse/dynwrap, version 0.1.0) and dynmethods (https://github.com/dynverse/dynmethods, version 0.1.0) to run the two best TI methods, found by the built-in TI method browser dynguidelines (https://github.com/dynverse/dynguidelines, version 1.0.1): PAGA-Tree (version 2.0.0) and Slingshot (version 2.0.1). All methods were run with default parameters and both methods take prior information of ground truth cluster (cell type) information and root cell information. Both dynwrap and dynmethods are part of the collection of R packages dynverse used in the manuscript by Saelens et al [1].

We use two evaluating metrics: Kendall’s *τ* and Hamming–Ipsen–Mikhailov distance (HIM). We designed a measure, *relative entropy*, to calculate how good the dataset is in terms of balancing the number of cells across all the cell states so that the underlined state tree is better preserved. Given a distribution of the states *X* = {*p*_1_, *p*_2_, *p*_3_, …, *p_N_* }, the relative entropy is calculated as following:

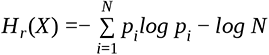

where 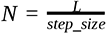 where *L* denotes the height of the cell state tree and *step_size* used to sample cell types on the state tree, and *p_i_* represents the frequency of the states at depth *i* on the state tree. The relative entropy subtracts the entropy of the uniform distribution of the same length so that the bias introduced by different *step_size* is removed. In other words, the relative entropy of a distribution is equal to the relative entropy of a piecewise constant interpolation of the distribution (Supplementary Note Sec. 2).

### Evaluation of ancestral gene expression inference methods

Using the simulated gene expression and lineage barcode data from the root to the leaf cells, we evaluate the accuracy of ancestral gene expression inference, performed by TreeVAE [40]. We compare TreeVAE with a VAE based baseline method. To take into account the accuracy of the lineage inference process, we either applied Cassiopeia [16] on the lineage barcodes of the leaf cells to obtain the reconstructed lineage or directly used the true lineage as the input.

We used TedSim to generate 10 datasets with 128 leaf cells and 126 internal cells. The gene expressions of the leaf cells are regarded as the input data for the algorithm and the gene expressions of the internal cells serve as the ground truth. We quantify the performances of TreeVAE and the baseline method in terms of three aspects between the inferred ancestral gene expression and the ground truth: 1. Cell to cell correlation(Spearman and Pearson correlation) and MSE (Mean Squared Error), using the true lineage(Fig. 7b). 2. Correlation and MSE of the average expression, using the reconstructed lineage and the true lineage (Supplementary Fig. 4).

### Evaluation of hybrid lineage inference methods on time-course data

TedSim is able to generate time-course scRNA-seq and lineage barcodes data by selecting cells at different depths of the lineage tree. Methods including LineageOT [46] and EntropicOT [8] were developed to infer the cell lineage relationships using time-course single cell gene expression data, where LineageOT also uses the cell lineage tree reconstructed from lineage tracing barcodes to improve their inference, thus is one of the “hybrid” methods we benchmark in this paper. We perform LineageOT in two different modes: using either the reconstructed cell lineage tree and using the true tree. The algorithms are tested on both continuous and discrete populations by tuning the parameters of our simulation (Fig. 7a).

We generate the dataset with a binary cell division tree of 512 leafs, and select two generations of cells to obtain time course data: the 512 descendant cells and 64 ancestor cells at the same depth. The ground truth of ancestor-descendant relationships is directly obtained from the cell division tree. We vary the *p_a_* and *step_size* to get both discrete and continuous trajectories: *p_a_* =0.4, *step_size* = 0.25 for the continuous dataset and *p_a_* =0.8, *step_size* = 0.75 for the discrete dataset. We benchmark the accuracy of the lineage inference using “ancestor prediction error” from the paper [18], which calculates the mean squared optimal transport distance between the inferred coupling matrix and the ground truth (Fig. 7a).

### Evaluation of lineage tree reconstruction methods

For this experiment, we use datasets generated by TedSim to compare the performances of lineage reconstruction methods: Cassiopeia (https://github.com/YosefLab/Cassiopeia, version 1.0.4), DCLEAR (https://github.com/ikwak2/DCLEAR, version 0.9.7) which use only the lineage barcodes and LinTIMaT (https://github.com/jessica1338/LinTIMaT), which also incorporates the gene expressions to refine cell lineage trees. We ran and compared two modes of Cassiopeia, Cassiopeia-greedy and Cassiopeia-hybrid, two modes of DCLEAR: DCLEAR-NJ and DCLEAR-FM and LinTIMaT. The detailed parameter settings to run the algorithms can be found in (Supplementary Note Sec. 4.2).

The datasets used to benchmark the lineage reconstruction methods vary in their mutation rate *μ*= (0.05, 0.1, 0.15, 0.2, 0.25, 0.3, 0.35, 0.4). We also compare their performances with and without genetic dropouts and under two different kinds of mutation event distribution: a synthetic uniform distribution and an experimental fitted distribution. For this experiment, we used the embryo2 from [14] to get the distribution of the experimental mutations, setting *N*_ms_ =100.

The reconstructed trees are compared to the same ground truth using two metrics: Robinson-Foulds distance and Triplet score. For the Robinson-Foulds distance, we use the API from the ete3 toolkit (https://doi.org/10.1093/molbev/msw046). For the Triplet score, we randomly select 10000 triplets and calculate the percentage of triplets that have the same structure in the reconstructed tree and the ground truth. To account for randomness in our simulation, we run the simulation 10 times with each combination of parameters and calculate the mean and standard deviation of the metrics.

## Acknowledgements

This work was supported in part by the US National Science Foundation DBI-2019771. Any opinions, findings and conclusions or recommendations expressed in this material are those of the authors and do not necessarily reflect the views of NSF.

## Author Contributions

X.P. and X.Z. conceived the study and designed the algorithm. X.P. implemented the package. X.P. and

H.L. performed the benchmarking analysis. X.P. and X.Z. drafted the manuscript. All authors revised the manuscript and approved the final manuscript.

---

**Supplementary Figure 1.**
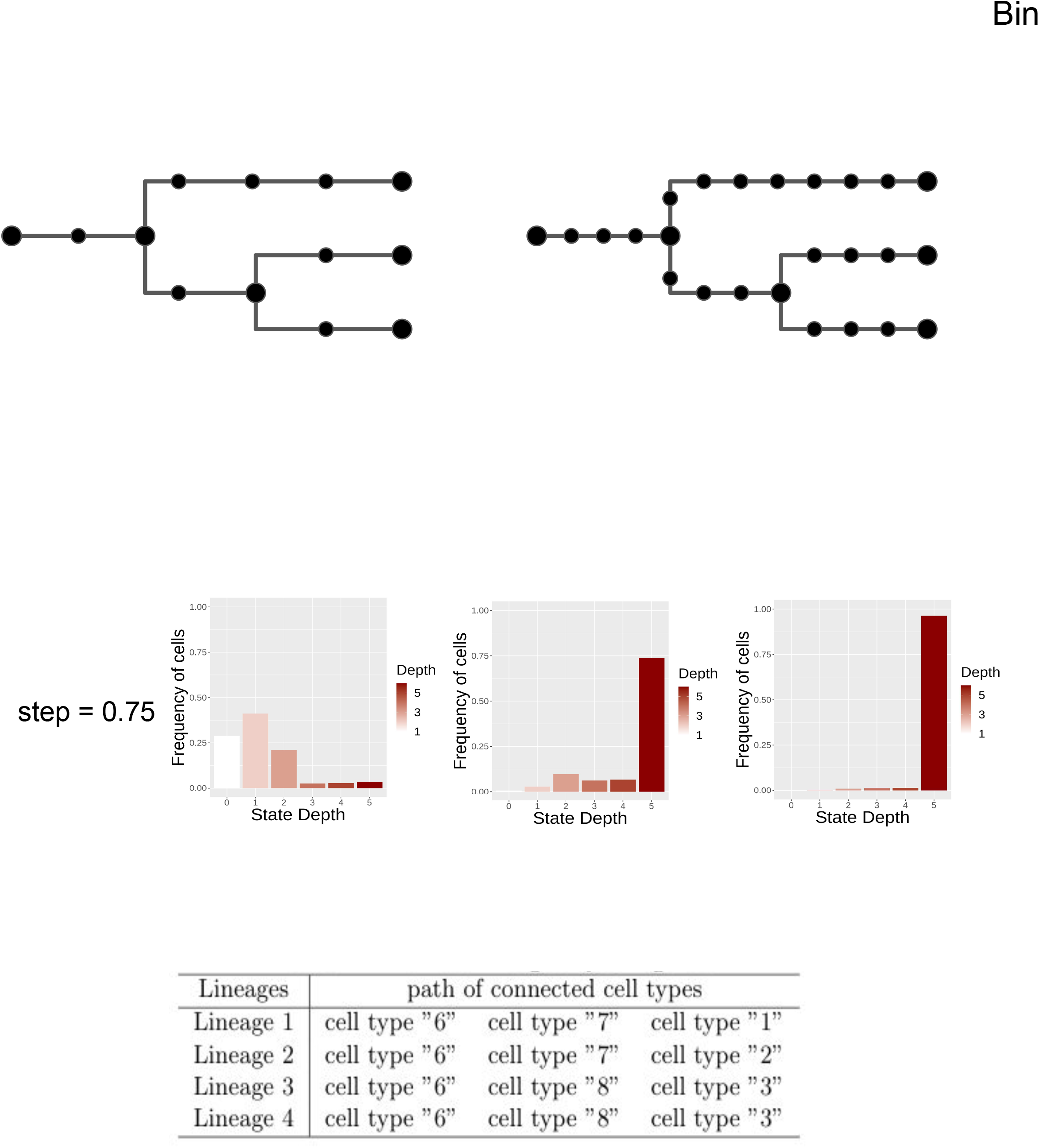
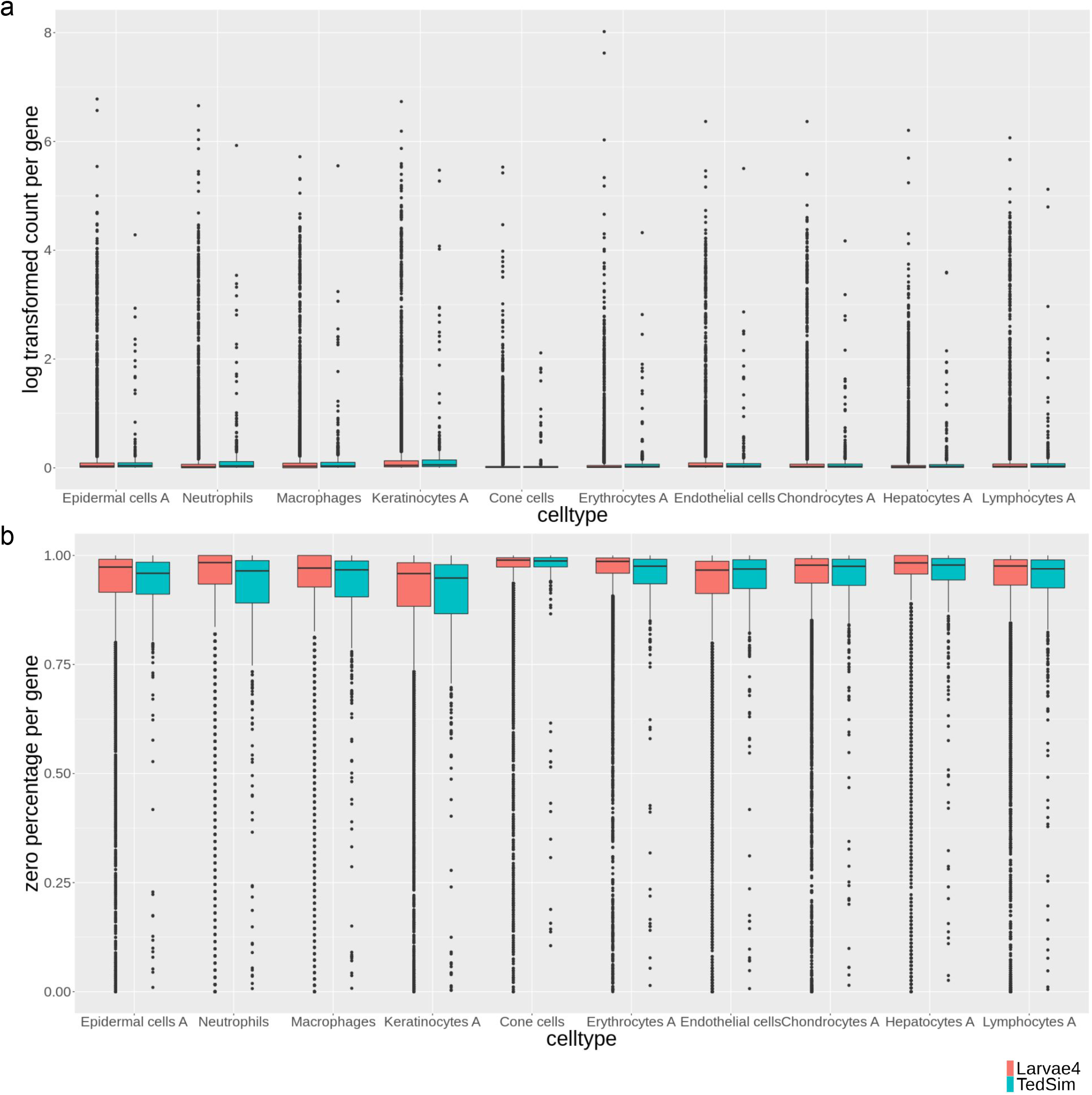
Statistical comparison between cell types of simulated data and gene expression of 5-dpf zebrafish larvae cells. Both metrics are calculated based on observed counts with capture efficiency *alpha_mean* = 0.1. (a) Average expression per gene. (b) Average zero percentages per gene.

**Supplementary Figure 2.**
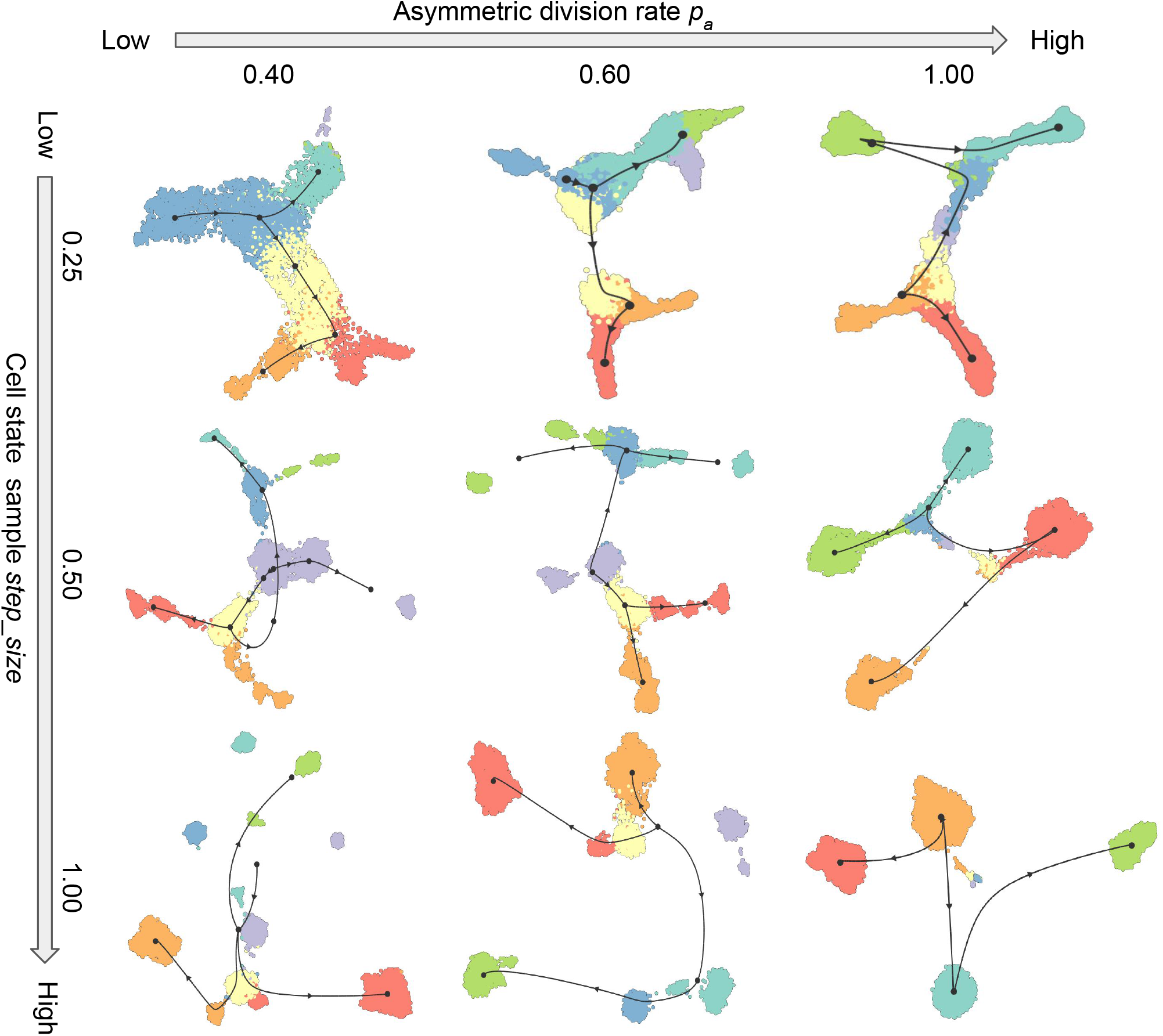
UMAP visualization of inferred trajectories by Slingshot with simulated gene expression data under various *p_a_* and *step_size* values. The TI methods are run with default settings from dynmethods (https://github.com/dynverse/dynmethods). The state tree used to generate the data can be referred to in Fig. 5a, and a randomly selected cell from cell type 6 is given as the root cell of the trajectories.

**Supplementary Figure 3.**
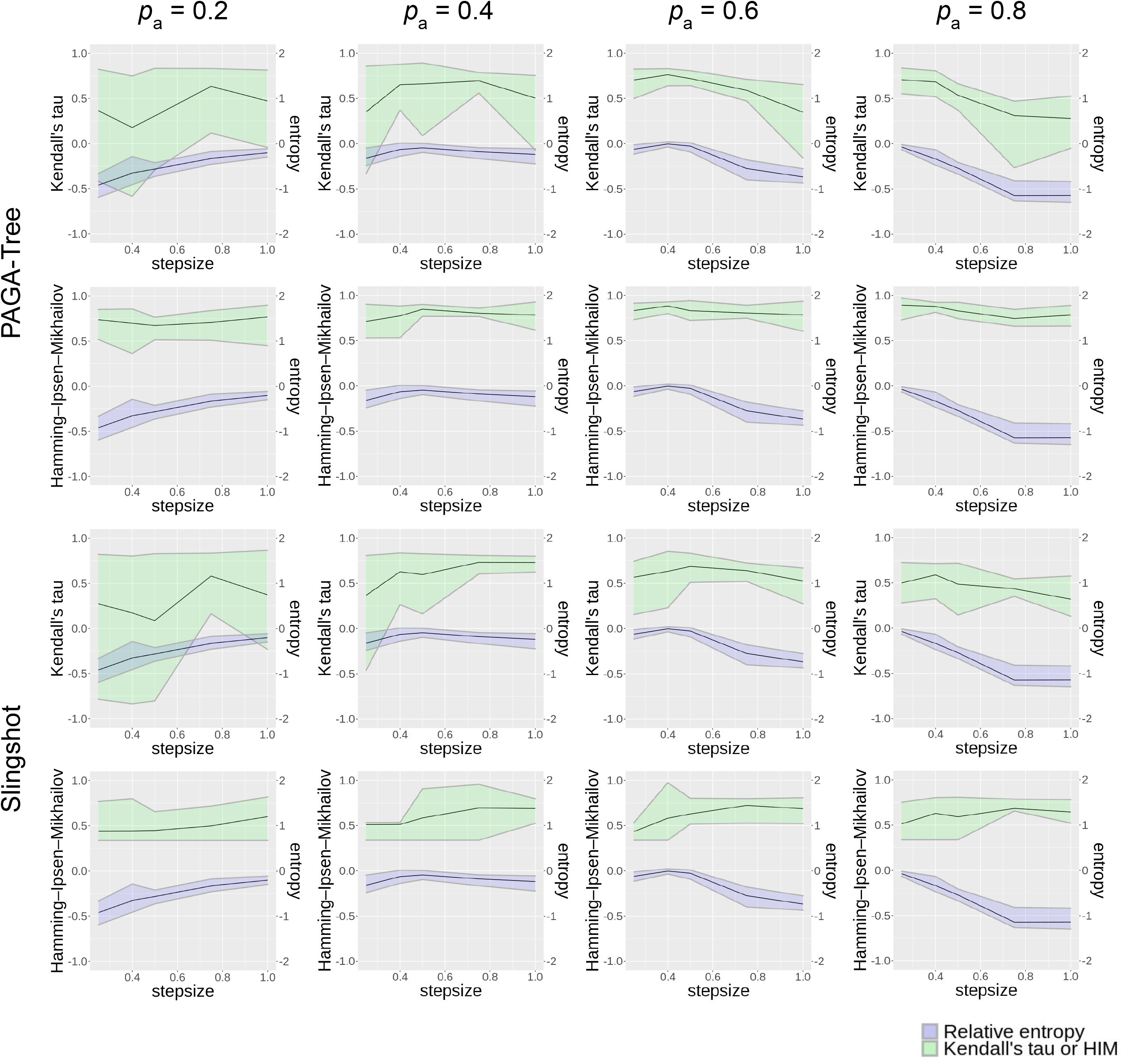
Pseudotime inference performances and relative entropy of the dataset with varying *step_size* parameters. The black curve represents the average score of 10 runs and the upper and lower bound of the ribbon represents maximum and minimum score. The PAGA-Tree results are the same as Fig. 6B. They are included here to facilitate the comparison between PAGA-Tree and Slingshot.

**Supplementary Figure 4.**
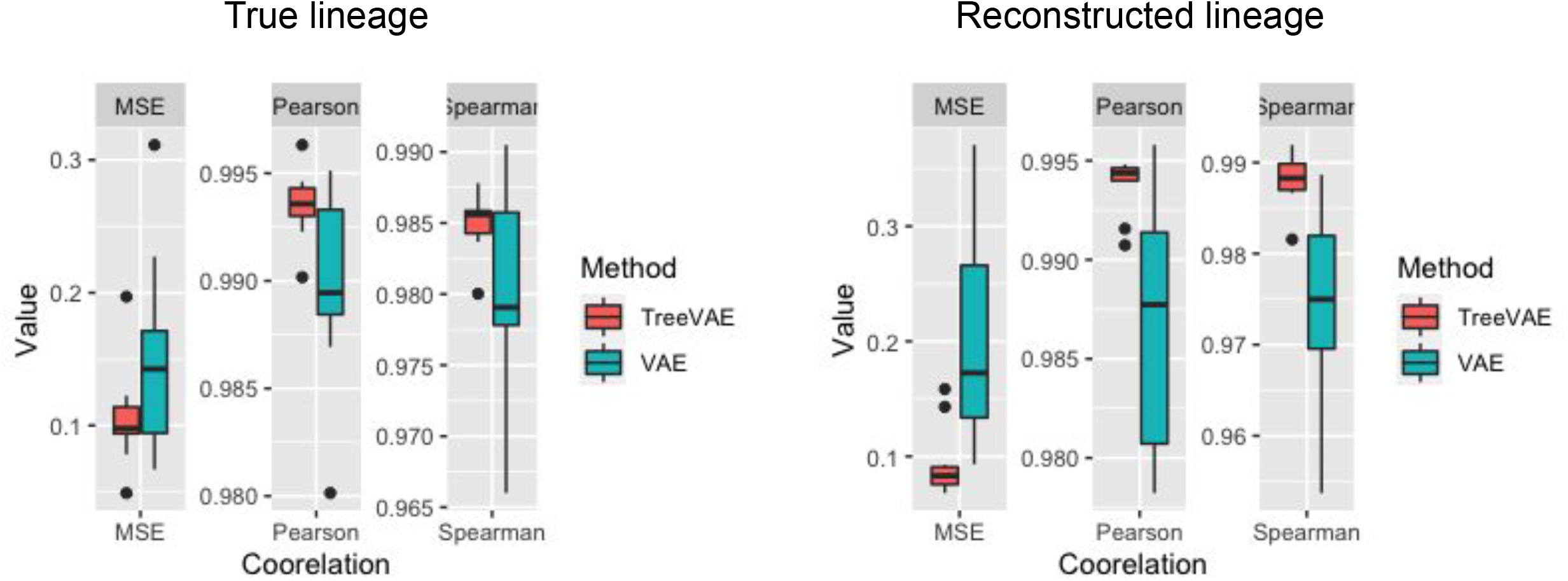
Correlation between average gene expression inferred and the ground truth. On the left, the input lineage is the true lineage used to generate the dataset while on the right, the input lineage is reconstructed using Cassiopeia. The average process is done because the number of internal nodes are different for the reconstructed lineage and the true lineage.

**Supplementary Figure 5.**
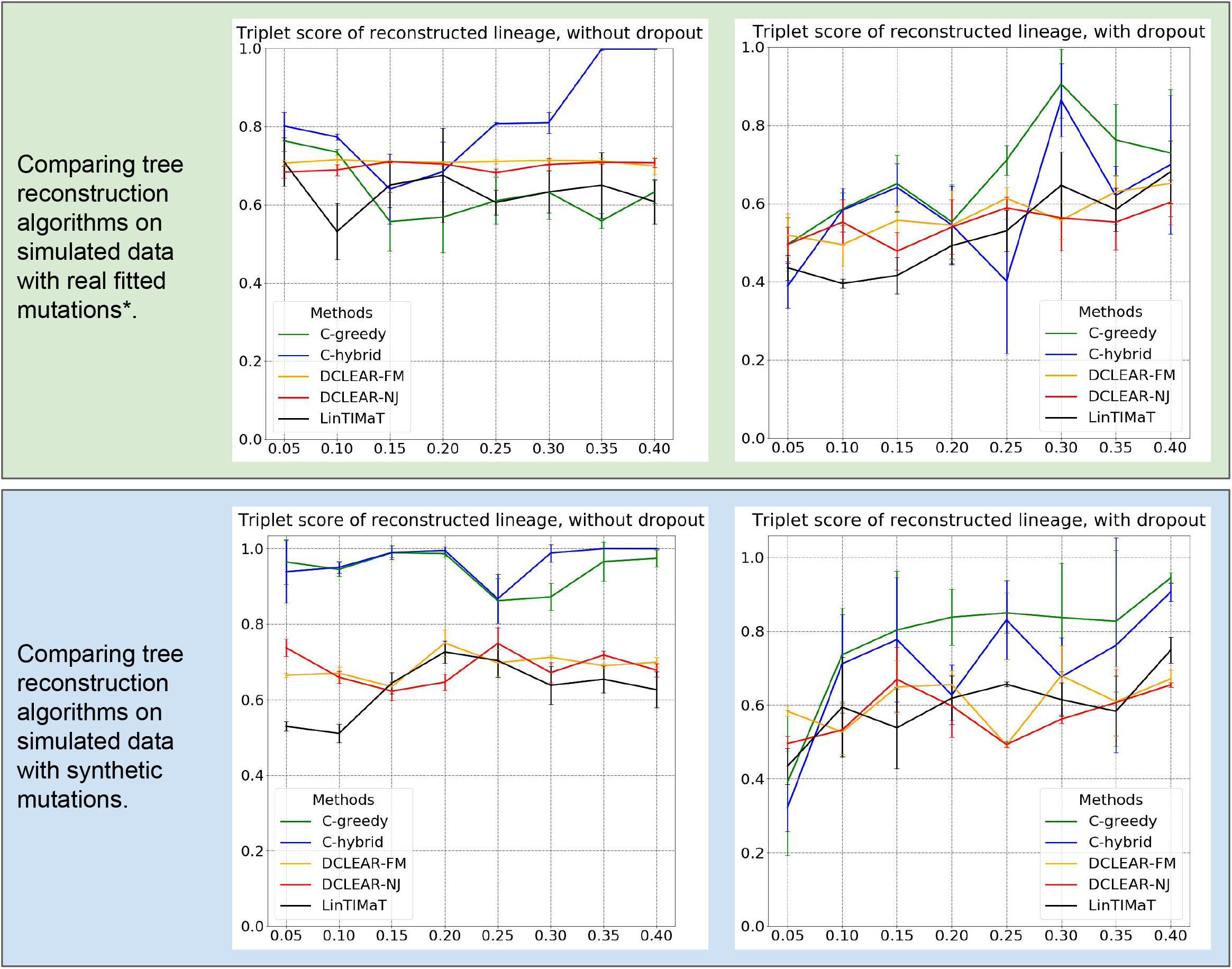
Benchmarking tree reconstruction methods on synthetic lineage barcodes using Triplet correct score. The mutation states are sampled from a fitted distribution of real mutations of embryo2 from M. Chan et al. (a) or a uniform distribution of synthetic mutations (b). Methods except LinTIMaT utilize only the lineage barcodes while LinTIMaT also uses gene expressions. Parameters used to simulate the datasets can be found in Supplementary Note Sec. 4.3. For each configuration (mutation rate and dropout), 10 simulations are run and used to test the methods.

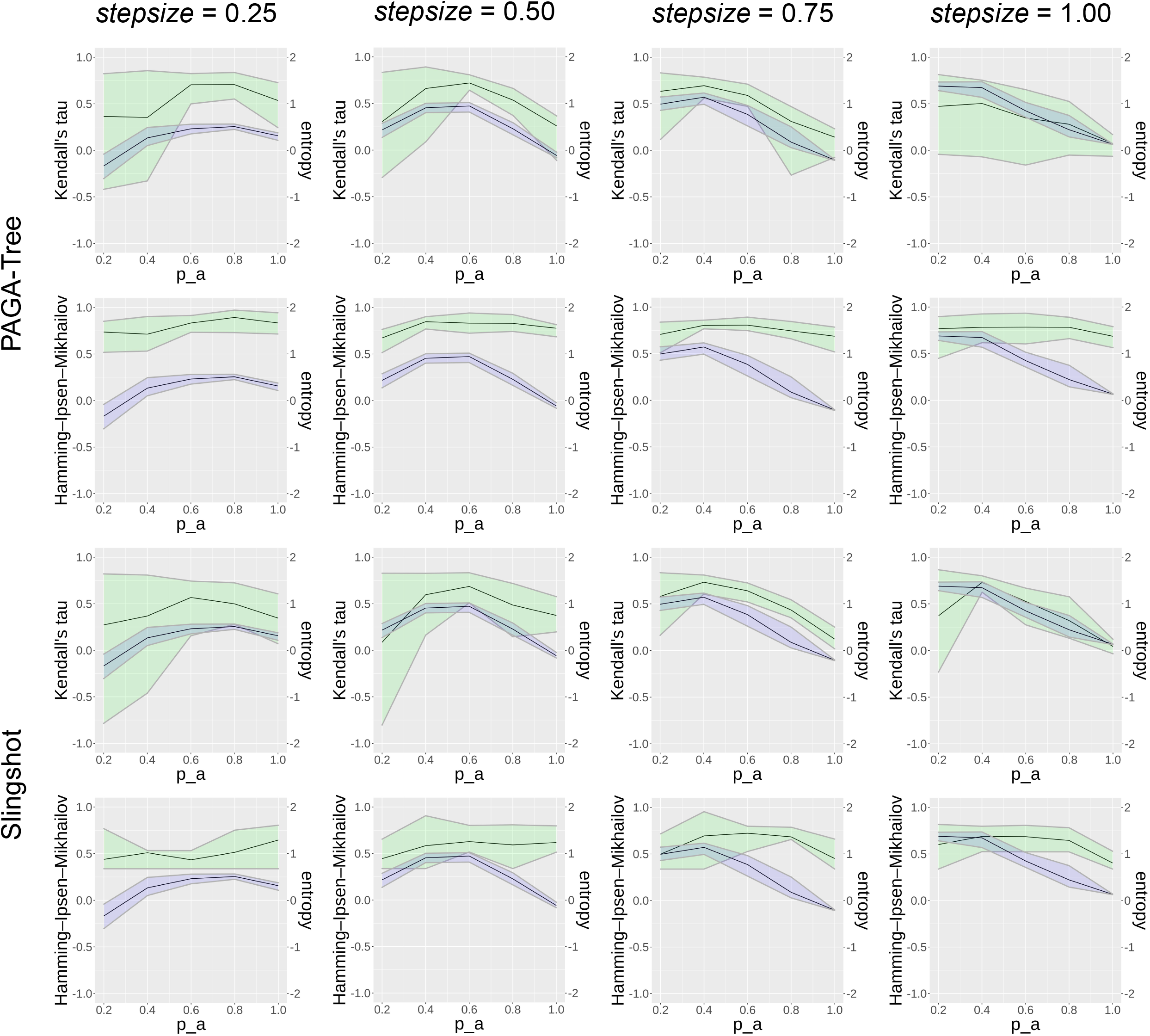

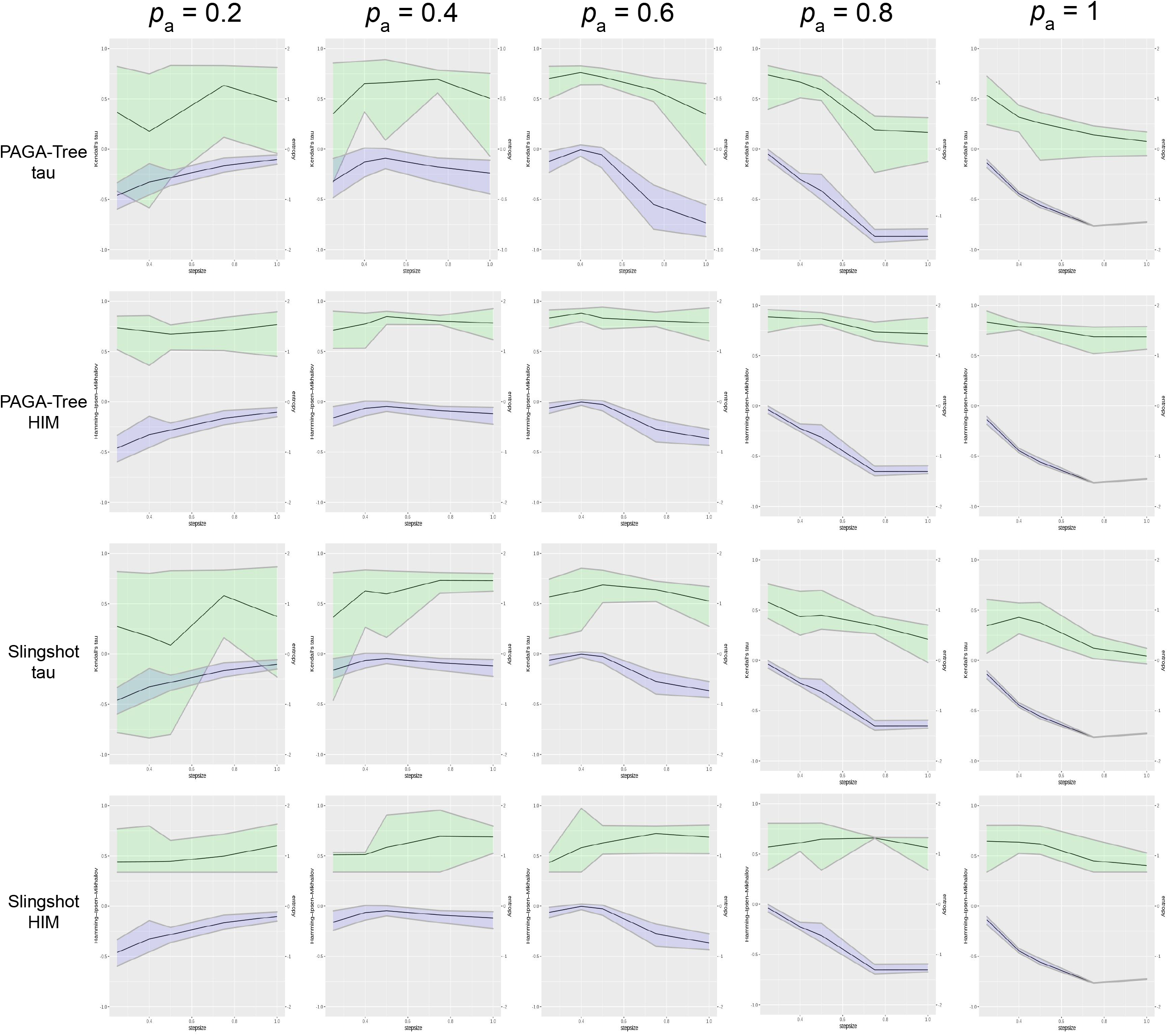

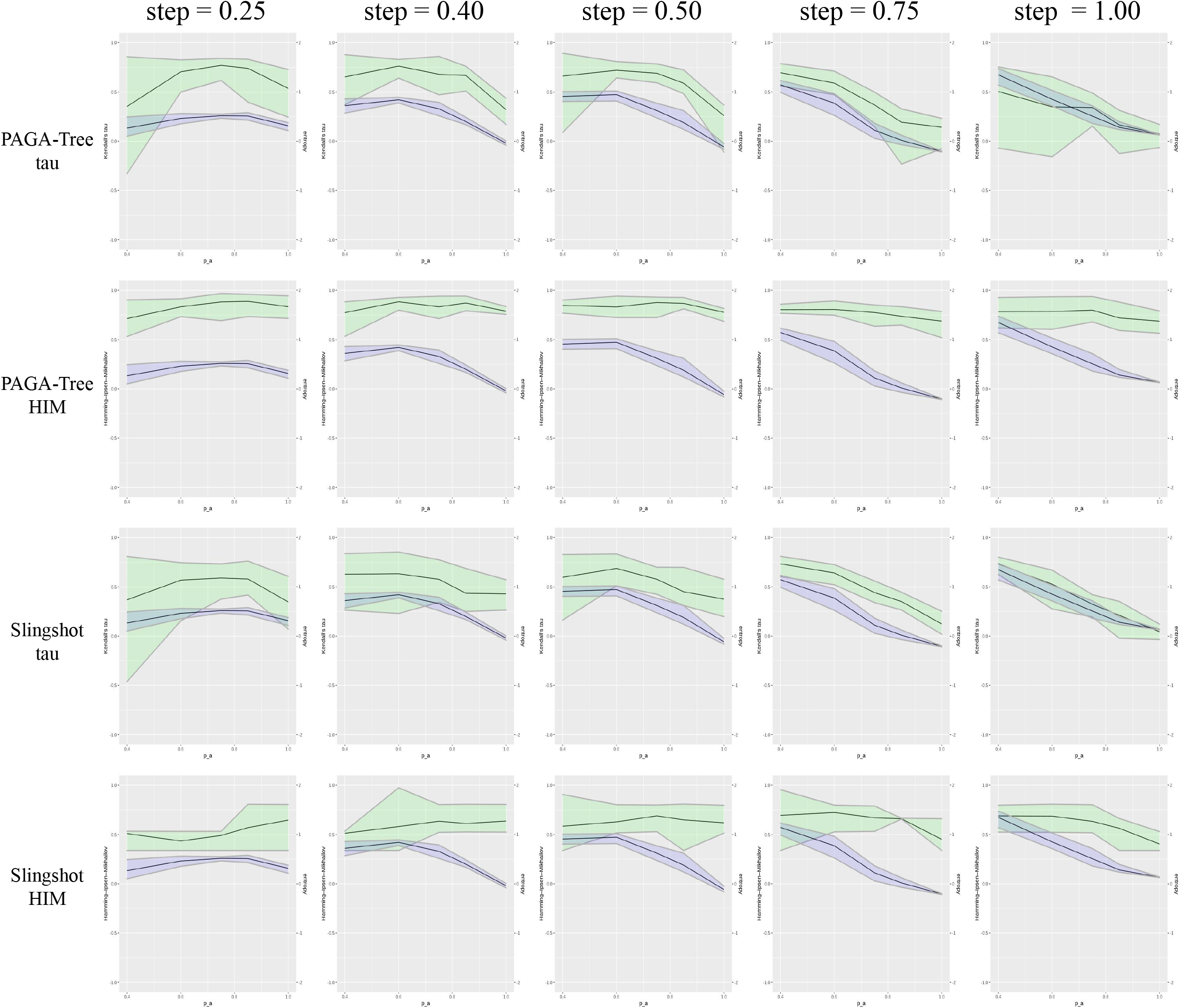

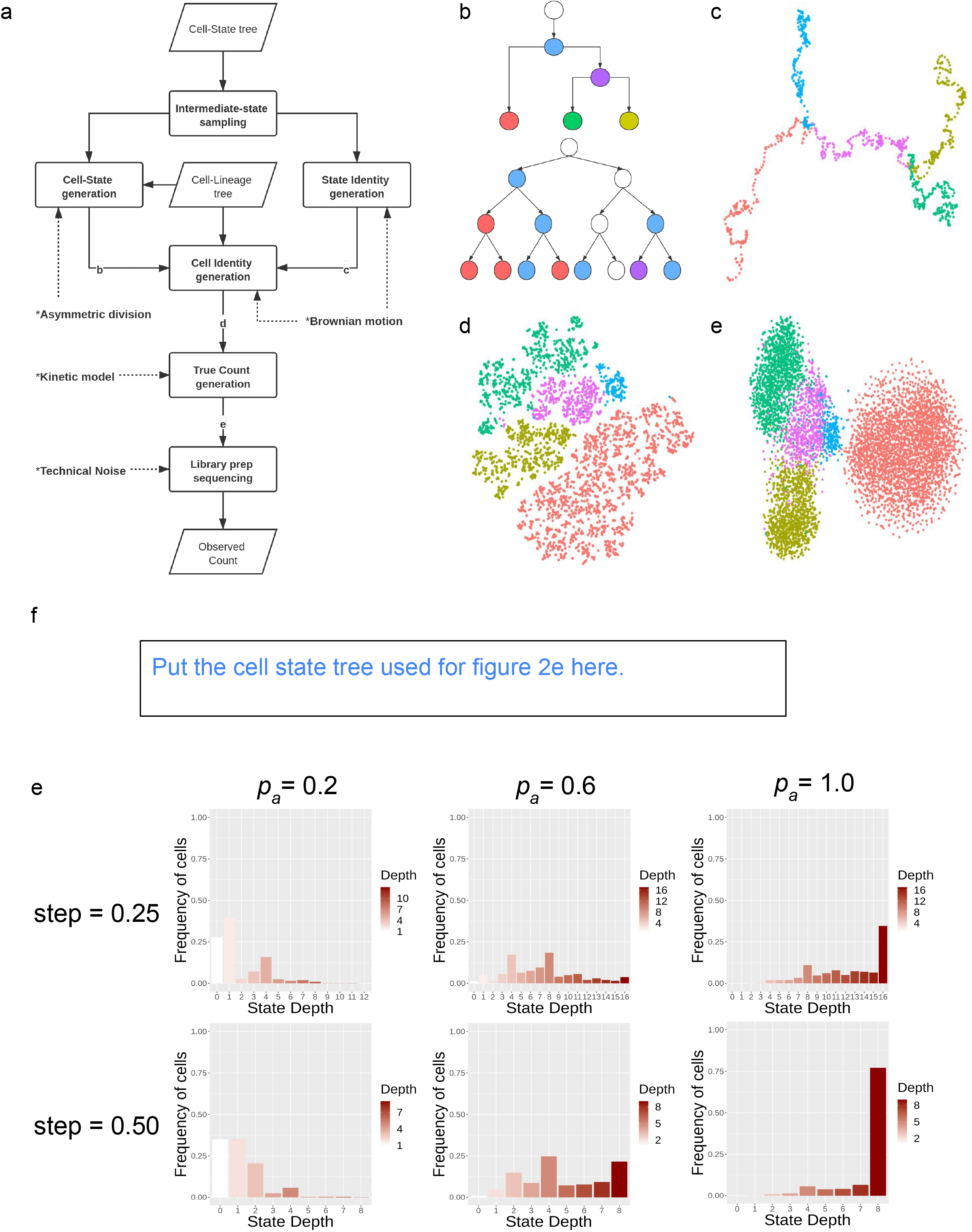

