## Supplementary Note for "TedSim: temporal dynamics simulation of single cell RNA-sequencing data and cell division history"

#### 1 Number of Cells (NoC) for one state path

Considering a cell division process of $k$ generations from one cell, there are $2^k$ leaf cells and $2^{k+1}-1$ cells in total. Considering a cell state path of $d$ states: $s_1, s_2, ..., s_d$ and asymmetric division rate $p_a$, and at each asymmetric division, one of the daughter cells will shift to the next state (+1), we can calculate the expected number of cells at each depth as following:

We denote the number of cells for state $s_i$ at the $j_{th}$ generation as $N_{s_i}(j)$. First, we write the recurrence relationship of number of cells at different states at the $j_{th}$ generation:
For $i = 1$

$$E\big[N_{s_i}(j)\big] = (2-p_a)N_{s_i}(j-1)$$

For $i = 2, 3, ..., d-1$

$$E\big[N_{s_i}(j)\big] = (2-p_a)N_{s_i}(j-1) + p_a N_{s_{i-1}}(j-1)$$

And for $i = d$

$$E[N_{s_d}(j)] = 2^j - \sum_{i=1}^{d-1} N_{s_i}(j) = 2N_{s_d}(j-1) + p_a N_{s_{d-1}}(j-1)$$

Clearly, the number of cells for the first state $s_1$ is only dependent on $p_a$, where higher $p_a$ results in less $s_1$ cells. For the middle states $s_2, s_3, ..., s_{d-1}$, the number of cells will be dependent on the previous state as well. Due to the finite number of total states, the end state will grow faster than all other states since it can only divide symmetrically, resulting in at least exponentially increasing at rate 2.

It is hard to come up with an analytical form for all the expectations for any specific $p_a$, but we are able to analyze a corner case, in which the asymmetric division rate $p_a = 1$.

Initial condition: $N_{s_1}(0) = 1$, $N_{s_i}(0) = 0$ for $i \geq 2$. At generation $k$, the expected number of cells can be given as following:

$$N_{s_1}(k) = N_{s_1}(k-1) = N_{s_1}(0) = 1$$

$$N_{s_2}(k) = N_{s_1}(k-1) + N_{s_2}(k-1) = 1 + N_{s_2}(k-1) = k$$

$$N_{s_3}(k) = N_{s_2}(k-1) + N_{s_3}(k-1) = k - 1 + N_{s_2}(k-1) = \frac{k(k-1)}{2}$$

...

In a more general way, if $d \leq k$, for $i = 1, 2, ..., d-1$

$$N_{s_i}(k) = \binom{k}{i-1}$$

And for $i = d$,

$$N_{s_d}(k) = 2^k - \sum_{i=1}^{d-1} \binom{k}{i-1} = \sum_{i=d}^{k+1} \binom{k}{i-1}$$

So that the distribution of the expected number of cells at different states are reduced to a binomial distribution for $p_a = 1$, and for $i < d$, the number of cells for $s_i$ increase at a polynomial speed while when $p_a < 1$, the speed will be at least exponential considering $E\big[N_{s_i}(j)\big] = (2-p_a)N_{s_i}(j-1) + p_a N_{s_{i-1}}(j-1) \geq= (2-p_a)N_{s_i}(j-1)$, so the number of cells at generation $k$ is at least $(2-p_a)^k$.

### 2 Relative Entropy unbiased for quantifying trajectory continuity

Given a probability mass function of a discrete random variable of N different possible outcomes:

$$p_N(X) = (p_1, p_2, p_3, ..., p_N)$$

The entropy of the variable:

$$H(p_N(X)) = -\sum_{i=1}^{N} p_i \log p_i$$

We define *relative entropy* $H_r(X)$ as:

$$H_r(X) = H(p_N(X)) - H(Unif(N)) = -\sum_{i=1}^{N} p_i \log p_i - \log N$$

where $Unif(N)$ denotes the discrete uniform distribution of length $N$. We claim that the relative entropy of a distribution is equal to the relative entropy of a piecewise constant interpolation of the distribution. Piecewise-constant interpolation of a distribution by factor $L$ (interpolating $L-1$ points between samples) can be defined as:

$$p_{LN}(X) = (p_1^1, p_1^2, ..., p_1^L, p_2^1, p_2^2, ..., p_2^L, ..., p_N^L)$$

where $p_i = \sum_{j=1}^{L} p_i^j$ (normalization) and $p_i^1 = p_i^2 = ... = p_i^L = p_i/L$ (piecewise-constant). Therefore:

$$\begin{aligned} H(p_{LN}(X)) &= -\sum_{i=1}^{N}\sum_{j=1}^{L} p_i^j \log p_i^j \\ &= -L\sum_{i=1}^{N} \frac{p_i}{L} \log \frac{p_i}{L} = \log L - \sum_{i=1}^{N} p_i \log p_i \\ &= \log L + H(p_N(X)) \end{aligned}$$

Which is the maximum entropy of $p_{LN}(X)$ after interpolation, which can be proven using Jensen's inequality. The relative entropy of $p_{LN}(X)$ is equal to $H_r(p_N(X)$:

$$\begin{aligned} H_r(p_{LN}(X)) &= H(p_{LN}(X)) - \log LN \\ &= H(p_N(X)) + \log L - \log LN \\ &= H(p_N X) - \log N \\ &= H_r(p_N(X)) \end{aligned}$$

In conclusion, the relative entropy measures how balanced the distribution

### 3 Proof of cell-cell distance reflecting lineage and state heterogeneity

#### 3.1 Tree diagrams

We use the cell state tree to model the cell differentiation process. The edge length models the distance between cell states. The cell lineage tree can be illustrated by a binary tree with uniform edge length.

(1) Cell State Tree:

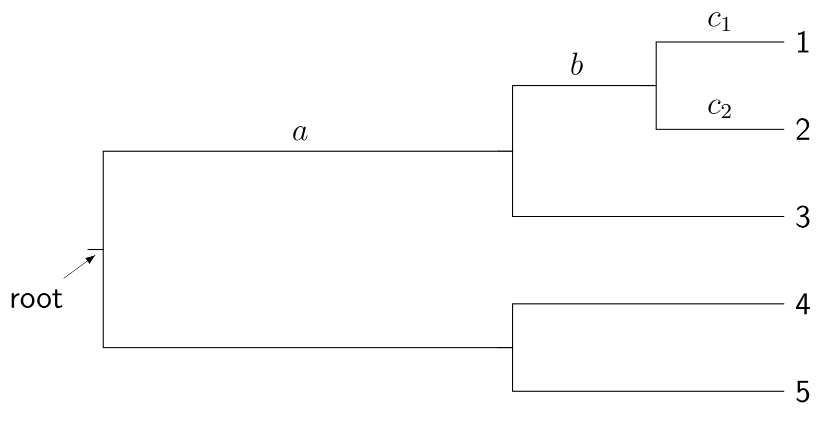


(2) Cell Lineage Tree:

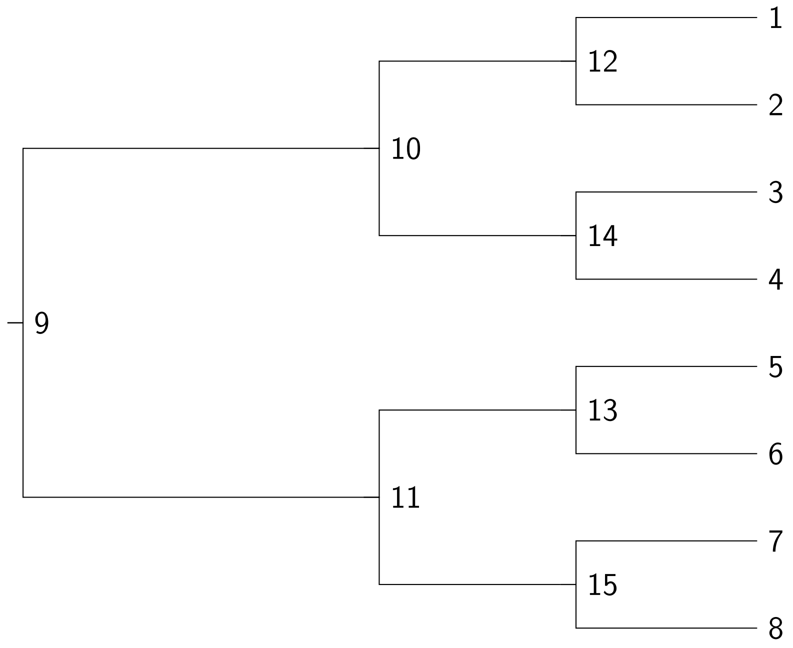


#### 3.2 Deriving trait differences of Brownian motion

(1) Trait values at the tips of the tree are normally distributed with mean $\mu$ and variance equal to the sum of branch lengths between the tips and the root, with $\mu$ being the value at the root.
Without loss of generosity, let $\mu = 0$:

$$W \sim \mathcal{N}(0, d)$$

We can reformulate Brownian Motion in terms of a 1 Dimensional Random Walk. In each unit time, the trait values take k steps of size $\frac{1}{\sqrt{k}}$ in random directions. Let the direction of the step be $\delta$ and it can take either value 1 or $-1$. The total distance of the random walk is:

$$W = \frac{1}{\sqrt{k}} \sum_{i=1}^{dk} \delta_i = \sqrt{d}(\frac{1}{\sqrt{dk}} \sum_{i=1}^{dk} \delta_i)$$

Because $\delta_i$ are i.i.d. with mean 0 and variance 1, by central limit theorem, their normalized sum $\sqrt{d}(\frac{1}{\sqrt{dk}} \sum_{i=1}^{dk} \delta_i)$ has standard normal distribution. Thus,

$$W \sim \mathcal{N}(0, d)$$

When the value at the root $\mu \neq 0$, we have $W = \mu + \frac{1}{\sqrt{k}} \sum_{i=1}^{dk} \delta_i \sim \mathcal{N}(\mu, d)$.

(2) Covariance of the trait values at the tips of the tree are equal to the branch length

between the root and their most recent common ancestor. Considering tip 1 and 2 on the state tree in **3.1**, where the trait value at the root is denoted as $\mu$:

$$W_1 \sim \mathcal{N}(\mu, a + b + c_1)$$

$$W_2 \sim \mathcal{N}(\mu, a + b + c_2)$$

Obviously, $W_1$ and $W_2$ are not independent.

$$\begin{aligned}
&Cov(W_1, W_2) = E(X - E(X))E(Y - E(Y)) \\
&= E(\mathcal{N}(0, a + b + c_1)\mathcal{N}(0, a + b + c_2)) \\
&= E((\mathcal{N}(0, a + b) + \mathcal{N}(0, c_1))(\mathcal{N}(0, a + b) + \mathcal{N}(0, c_2))) \\
&= E(\mathcal{N}(0, a + b)\mathcal{N}(0, a + b) + \mathcal{N}(0, a + b)\mathcal{N}(0, c_1) + \mathcal{N}(0, a + b)\mathcal{N}(0, c_2) + \mathcal{N}(0, c_1)\mathcal{N}(0, c_2)) \\
&= E(\mathcal{N}^2(0, a + b)) = Var(\mathcal{N}(0, a + b)) \\
&= a + b
\end{aligned}$$

where we utilize the fact that the expectation between two independent zero-mean random variables is zero. The result indicates that the covariance of the trait values are equal to the shared branch length between two tips.

(3) Mean-shifting Brownian motion captures both cell state and cell lineage.

In TedSim, we first perform Brownian motion on the state tree to get the State Identity Factors (SIFs), and then we perform Brownian motion on the lineage tree, and at the same time add the specific state means given the state of the cell and the SIFs.
Considering two tips on the cell lineage tree in **3.1**, cell 5 and cell 7, the trait values on the cell lineage tree can be calculated as following (the edges of the cell lineage tree are considered to be 1):

$$F_5 = W_{state(5)} + (W_{9-11} + W_{11-13} + W_{13-5})$$

$$F_7 = W_{state(7)} + (W_{9-11} + W_{11-15} + W_{15-7})$$

Similarly to (2), we can calculate the covariance between cell 5 and cell 7 as:

$$\begin{aligned}
Cov(F_5, F_7) =& E((W_{state(5)} + (W_{9-11} + W_{11-13} + W_{13-5}))(W_{state(7)} + (W_{9-11} + W_{11-15} + W_{15-7}))) \\
&- E(W_{state(3)} + (W_{9-11} + W_{11-13} + W_{13-5}))E(W_{state(7)} + (W_{9-11} + W_{11-15} + W_{15-7})) \\
&= (E(W_{state(5)}W_{state(7)}) - E(W_{state(5)})E(W_{state(7)})) + E(W_{9-11}^2) - E^2(W_{9-11}) \\
&= Cov(W_{state(5)}, W_{state(7)}) + Var(W_{9-11})
\end{aligned}$$

which indicates that the covariance between two cells consists of two parts: the covariance between the states of the cells and the variance of the shared ancestors of the cells. The first term can be further translated as the shared hidden states on the cell state tree.

### 4 Simulation settings

#### 4.1 TedSim simulations for fitting real datasets

(1) State tree inferred by hierarchical clustering:

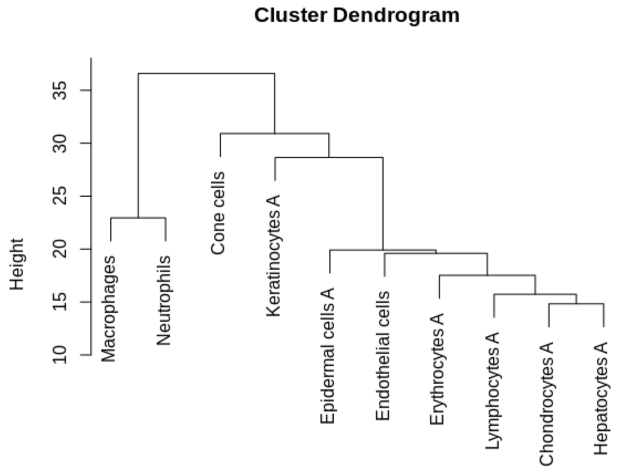


Figure 1: Hierarchical clustering on the mean expressions of the cell types

The edge lengths are then rounded up to the nearest integer so that at least one state is sampled for each edge.

(2) State-varying parameters:

In order to achieve state-varying average counts and zero percentages, we tune the $scale\_s$ parameter to reflect the corresponding UMI counts per cell type in the real dataset:

$$scale\_s = 0.03UMI\_mean$$

(3) Other Simulation parameters:

| | |
|---|---|
| Number of cells (ncells) | 1024 |
| Number of genes (ngenes) | 500 |
| Number of Identity Factors ($N_{IF}$) | 30 |
| Identity Factor center (starting value for diff-IF) | 1 |
| Number of diff-Identity Factors ($N_{diff}$) | 20 |
| nondiff-SIF standard deviation ($\sigma$) | 0.5 |
| Probability of nonzero gene effect (ge_prob) | 0.3 |
| Probability of outlier gene (prob_hge) | 0.03 |
| Mean of capture efficiency $\alpha$ (alpha_mean) | 0.1 |
| Standard deviation of capture efficiency $\alpha$ (alpha_sd) | 0.1 |

#### 4.2 TedSim simulations for trajectory inference methods

(1) State Tree:

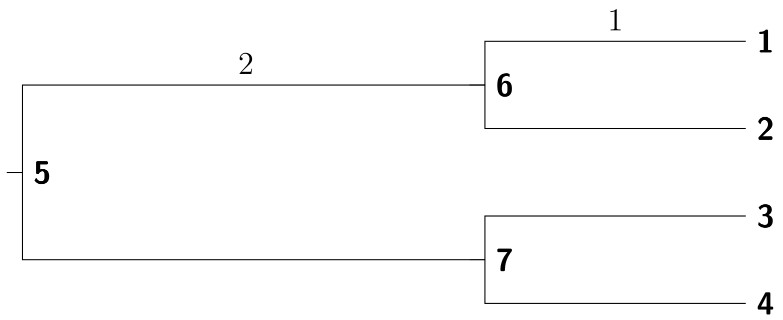


(2) Variables: we run 10 instances for each combination of the variables.
Asymmetric division rate: $p_a = [0.2\ 0.4\ 0.6\ 0.8\ 1.0]$.
State Identity Factor stepsize: $step = [0.2\ 0.25\ 0.5\ 0.75\ 1]$.
(3) Other Simulation parameters:

| | |
|---|---|
| Number of cells (ncells) | 8192 |
| Number of genes (ngenes) | 500 |
| Number of leaf states (n_leaf_states) | 4 |
| Number of Identity Factors ($N_{IF}$) | 30 |
| Identity Factor center (starting value for diff-IF) | 1 |
| Number of diff-Identity Factors ($N_{diff}$) | 20 |
| nondiff-SIF standard deviation ($\sigma$) | 0.5 |
| Probability of nonzero gene effect (ge_prob) | 0.3 |
| Probability of outlier gene (prob_hge) | 0.015 |
| Mean of capture efficiency $\alpha$ (alpha_mean) | 0.2 |
| Standard deviation of capture efficiency $\alpha$ (alpha_sd) | 0.05 |

##### 4.3 TedSim simulations for lineage reconstruction algorithms

(1) State Tree:

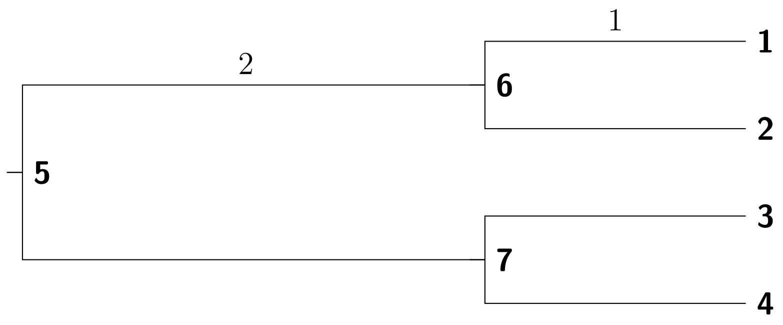


(2) Variables:

Mutation rate: $\mu = [0.05,0.1,0.15,0.2,0.25,0.3,0.35,0.4]$.

Dropout: $p_d = 0$ or 1.

Distribution of mutated states: $unif_{on} = 0$ or 1.

We run 10 instances for each combination of the variables.

(3) Other Simulation parameters:

| | |
|---|---|
| Number of cells (ncells) | 512 |
| Number of genes (ngenes) | 500 |
| Number of leaf states (n_leaf_states) | 4 |
| Asymmetric division rate ($p_a$) | 0.4 |
| State Identity Factor stepsize ($step$) | 0.25 |
| Number of Identity Factors ($N_{IF}$) | 30 |
| Identity Factor center (starting value for diff-IF) | 1 |
| Number of diff-Identity Factors ($N_{diff}$) | 20 |
| nondiff-SIF standard deviation ($\sigma$) | 0.5 |
| Probability of nonzero gene effect (ge_prob) | 0.3 |
| Probability of outlier gene (prob_hge) | 0.015 |
| Mean of capture efficiency $\alpha$ (alpha_mean) | 0.2 |
| Standard deviation of capture efficiency $\alpha$ (alpha_sd) | 0.05 |
| Number of target sites (n_char) | 32 |
| Number of possible mutated states (N_ms) | 100 |

(4) Lineage reconstruction methods settings:

LinTIMaT: number of genes: -gc 100, number of mutation likelihood iterations: -mi 100000, number of combined likelihood iterations: -ci 50000.

DCLEAR: Dropout weight: 12

Cassiopeia-hybrid: time limit for one ILP instance: 3600, maximum neighborhood size for ILP instances: 10000, cell neighborhood cutoff: 200.

##### 4.4 TedSim simulations for integrative lineage inference methods

### 5 Pseudocode for simulating Cell identity Factors and Barcodes

**Algorithm 1** State Identity Factors(SIF) Generation

```
function SIFGenerate(T_state, n_diff, resolution)
    Initialize Node_par as the root of T_state
    Initialize Ψ
    function SampleSIF(Node_par,T_state,n_diff,Ψ,resolution)
        for every child Node_child of Node_par on T_state, do
            Find the edge (Node_par,Node_child) on T_state
            for i = 1, 2, ..., n_diff do
                compute Brownian motion distance d for all states on edge,
                given the edge and resolution of the random walk
                Ψ ← Ψ ∪ [Node_par, Node_child, depth, d]
            end for
            Ψ ← SampleSIF(Node_child,T_state,n_diff,Ψ)
        end for
        return Ψ
    end function
    return Ψ
end function
```

**Algorithm 2** Simulating States for all cells

**function** SimulateCellStates($Node_{par}$,$T_{state}$,$T_{cell}$,$S$,$\Psi$,$p_a$)
    Get the $Node_{child}$ of $Node_{par}$ on $T_{cell}$
    Get the cell state of $Node_{par}$: $state_{par} \leftarrow S[Node_{par}]$
    Select one of the child $Node_{child1}$ of $Node_{par}$ on $T_{cell}$
    **for** every child $Node_{child}$ of $Node_{par}$ on $T_{cell}$, **do**
        $S(Node_{child}) \leftarrow state_{par}$
        **if** $state_{par}$ is not a leaf state **then**
            **if** asymmetric division: $U(0,1) \leq p_a$ **then**
                **if** $Node_{child} == Node_{child1}$ **then**
                    $S(Node_{child}) \leftarrow$ 'Next' state from $state_{par}$
                **end if**
            **end if**
        **end if**
        $S \leftarrow$ SimulateCellStates($Node_{child}$,$T_{state}$,$T_{cell}$,$S$,$\Psi$,$p_a$)
    **end for**
    **output** $S$
**end function**

**Algorithm 3** Generate Mutated barcode

**function** GenerateMutation($b$)
    $b_{out} \leftarrow b$
    Generate synthetic mutated states $states_m$ with distribution $p$
    With mutation rate $\mu$, find mutation sites mu_site ←Unif(where $b \neq 0$)$\leq \mu$
    Mutate the selected sites: $b_{out}[mu\_loc]$ ←sample($states_m$,$p$)
    **if** length(mu_site)$\geq 2$ **then**
        Randomly select two sites (a,b) from mu_site
        Drop the characters in between: $b_{out}[a:b]$ ←'-'
    **end if**
    **output** $b_{out}$
**end function**

**Algorithm 4** Simulating Cell Identity Factors (CIFs) and Barcodes

**input** $N_{cell}, T_{state}, p_a, n_{CIF}, n_{diff}, resolution$
Generate a bifurcating tree $T_{cell}$ of $N_{cells}$ tips
Get the total number of nodes $N_{nodes}$ of $T_{cell}$
$\Psi \leftarrow$ SIFGENERATE($T_{state}$, $n_{diff}$, $resolution$)
$S \leftarrow$SIMULATECELLSTATES($Node_{par}$,$T_{state}$,$T_{cell}$,$S$,$\Psi$,$p_a$)
**Initialize** $Node_{par}$ as the root of $T_{cell}$
**Initialize** Barcode matrix $M$ as zero matrix (unmutated)
**Initialize** Cell Identity Factors matrix $CIF$
Generate non-Diff CIF values for all cells $CIF(nondiff) \leftarrow N(1, 0)$
**function** SAMPLELINEAGE($Node_{par}$,$\Psi$,$CIF$, $S$, $M$)
    Get the diffCIFs of $Node_{par}$: $CIF_{par} \leftarrow CIF(Node_{par})(diff)$
    Get the parent barcode: $M_{par}$ from $M$
    Get the parent state: $state_{par} \leftarrow S[Node_{par}]$
    **for** every child $Node_{child}$ of $Node_{par}$ on $T_{cell}$, **do**
        $CIF(Node_{child})(diff) \leftarrow CIF_{par}$
        Get the current cell state: $state \leftarrow S[Node_{child}]$
        $CIF(Node_{child})(diff) += \Psi(state, 4) - \Psi(state_{par}, 4) + N(0, 1)$
        $M_{child} \leftarrow$GENERATEMUTATION($M_{par}$), and update $M$
        $CIF, M \leftarrow$SAMPLELINEAGE($Node_{child}$,$\Psi$,$CIF$,$S$,$M$)
    **end for**
    **return** $CIF, M$
**end function**
Leave out internal nodes in $CIF$,$M$ and $S$
**output** $T_{cell}$,$CIF$,$S$,$M$

**Algorithm 5** Get the 'Next' state on the tree

**function** WALKTREE($s_{par}$,$step_w$)
    $b_{out} \leftarrow b$
    Generate synthetic mutated states $states_m$ with distribution $p$
    With mutation rate $\mu$, find mutation sites mu_site $\leftarrow$Unif(where $b \neq 0$)$\leq \mu$
    Mutate the selected sites: $b_{out}[mu_loc] \leftarrow$sample($states_m$,$p$)
    **if** length(mu_site)$\geq$ 2 **then**
        Randomly select two sites (a,b) from mu_site
        Drop the characters in between: $b_{out}[a : b] \leftarrow$'-'
    **end if**
    **output** $b_{out}$
**end function**
